# Examining the Biophysical Properties of the Inner Membrane of Gram-Negative ESKAPE Pathogens

**DOI:** 10.1101/2024.08.13.607768

**Authors:** Golbarg Gazerani, Lesley R. Piercey, Syeda Reema, Katie A. Wilson

**Affiliations:** Department of Biochemistry, Memorial University of Newfoundland, St. John’s, Newfoundland and Labrador, Canada

**Author notes:** These authors contributed equally (Golbarg Gazerani, Lesley R. Piercey).

## Abstract

The World Health Organization has identified multidrug-resistant bacteria as a serious global health threat. Gram-negative bacteria are particularly prone to antibiotic resistance, and their high rate of antibiotic resistance has been suggested to be related to the complex structure of their cell membrane. The outer membrane of Gram-negative bacteria contains lipopolysaccharides that protect the bacteria against threats such as antibiotics, while the inner membrane houses 20-30% of the bacterial cellular proteins. Given the cell membrane’s critical role in bacterial survival, antibiotics targeting the cell membrane have been proposed to combat bacterial infections. However, a deeper understanding of the biophysical properties of the bacterial cell membrane is crucial for developing effective and specific antibiotics. In this study, Martini coarse-grain molecular dynamics simulations were used to investigate the interplay between membrane composition and biophysical properties of the inner membrane across four pathogenic bacterial species: *Klebsiella pneumoniae, Pseudomonas aeruginosa, Enterobacter cloacae*, and *Escherichia coli*. The simulations indicate the impact of species-specific membrane composition on dictating the overall membrane properties. Specifically, cardiolipin concentration in the inner membrane is a key factor influencing the membrane features. Model membranes with varying concentrations of bacterial lipids (phosphatidylglycerol, phosphatidylethanolamine, and cardiolipin) further support the significant role of cardiolipin in determining the membrane biophysical properties. The bacterial inner membrane models developed in this work pave the way for future simulations of bacterial membrane proteins and for simulations investigating novel strategies aimed at disrupting the bacterial membrane to treat antibiotic-resistant infections.

## Introduction

Antibiotic resistance is a global health emergency. In 2019, it was estimated that 4.95 million people died worldwide due to an antibiotic resistant bacterial infection.^1^ Furthermore, without the creation of new strategies to combat antibiotic resistant infections, bacterial infections that were once easily treatable will once again lead to death. It is estimated that at the current rate, antibiotic resistant infections could kill 10 million people a year by 2050.^2, 3^ The ESKAPE bacterial pathogens (*Enterococcus faecium, Staphylococcus aureus, Klebsiella pneumoniae, Acinetobacter baumannii, Pseudomonas aeruginosa,* and *Enterobacter spp.*) are the bacteria species that are the leading cause of life-threatening nosocomial infections around the world.^4^ The high rate of infection is due to the highly virulent and multi-drug resistant nature of these pathogens that allow them to “escape” commonly used antibiotics.^4^ Furthermore, the ESKAPE pathogens are included on the 2017 and 2024 World Health Organization (WHO) priority pathogen lists, which identifies pathogens that pose the greatest threat to human health and therefore are a priority for research and the development of new treatments.^5, 6^ Among these pathogens, *A.baumannii and* Enterobacterales (e.g., *Enterobacter spp*. and *Escherichia coli*) have the highest priority (Priority 1 or critical priority) on the 2024 WHO priority pathogen list, while *P.aeruginosa, S.aureus* and *E.faecium* are categorized by the WHO as Priority 2 or high priority. It is noteworthy that all priority 1 pathogens identified by the WHO are Gram-negative bacteria.

Gram-negative bacteria have a distinctive cell envelope that is comprised of two bilayers, namely the inner membrane and the outer membrane. While the outer membrane is characterized by lipopolysaccharides in the outer leaflet and phospholipids in the inner leaflet, the inner membrane is composed of phospholipids in both leaflets. Key phospholipids found in Gram-negative bacterial cell membranes are phosphatidylethanolamine (PE), phosphatidylglycerol (PG) and cardiolipin (CL).^7^ These lipids all exhibit variation in the lipid tail length and degree of unsaturation. Typical lipid tails within the Gram-negative bacterial cell membrane contain between 14 and 18 carbons, arranged as straight chains or containing cyclopropane groups.^8^ The cell envelope plays an important role in protecting the Gram-negative bacteria from threats in their environment. Beyond providing a protective barrier around the cell and creating an enclosed environment that enables cellular life, the cell membranes contain numerous proteins critical for cellular functions. For example, in *E.coli*, ∼20-30% of the cellular proteins are found in the inner membrane and ∼2% are found in the outer membrane.^9^ The proteins found within the inner membrane are involved in diverse functions such as energy production, lipid biosynthesis, protein secretion, and transport.^9^ Furthermore, the protein-lipid interactions play an important role in modulating the activity of membrane proteins.^10^ Due to the cell envelope’s critical role in essential process for bacteria, antibiotics that target the cell membrane, such as antimicrobial peptides, have been proposed to combat antibiotic resistant infections.^11, 12^ Nevertheless, gaining deeper insights into the biophysical properties of the bacterial cell membrane is imperative for developing effective and specific antibiotics targeting these infections.

A common method used to understand the biophysical properties of membranes is molecular dynamics (MD) simulations.^13–15^ Long-timescale MD simulations of realistic membrane environments is an excellent way to gain insight into the properties of bacterial cell membranes and assess the impact of individual lipid components on membrane biophysical properties.^16^ However, historically membranes have been modeled with simplified compositions for short timescales.^14^ Specifically, as PE and PG are the most common lipid classes found in the bacterial cell envelope, traditional simulation of bacterial membranes often only included two lipid species such as POPE and POPG,^17^ while other studies model bacterial membranes with a single lipid species (e.g., POPE).^18^ In these models the impacts of variations in lipid tail composition and the impact of minor lipid species such as CL were overlooked. Recent studies emphasize the necessity of adopting realistic lipid compositions to accurately represent membrane biophysics. For instance, coarse-grain investigations into the *E.coli* inner membrane reveal that a simplistic model composed of three lipid species fails to replicate the properties of a complex, biologically relevant membrane containing 14 lipid species.^19^ Furthermore, introducing cyclopropane fatty acids into 14-component coarse-grain *E.coli* inner membrane^20^ or into a 4-7 component atomistic PE/PG membrane^21^ reduces the membrane packing density and thereby enhances membrane fluidity.^20^ While, atomistic simulations of 5-8 component PE/PG membranes representing *P.aeruginosa* in the planktonic or biofilm mode show that increased lipid tail length and increasing PG lipid concentration lead to a less rigid and thicker membrane.^22^ Finally, increasing lipid tail unsaturation in a coarse-grain model of the *A.baumannii* inner membrane with complex lipid compositions (10-13 lipid species) leads to increases in lateral lipid diffusion and area per lipid, as well as modulation of domain formation.^23^ Collectively, the previous work underscores the impact of lipid composition on the bacterial inner membrane biophysical properties and therefore highlights the importance of studying bacterial membranes with realistic lipid compositions.

To the best of our knowledge *E.coli*^19, 20^ and *A.baumannii*^23^ are the only Gram-negative bacterial species that the inner membrane has been computationally studied with a realistic lipid composition. These two species vary in the concretion of lipid classes (5% CL, 22% PG, 73% PE for *E.coli* and 28% CL, 48% PG, 24% PE for *A.baumannii*) and tail group saturation (i.e., *E.coli* membrane contains cyclopropane fatty acids and 0.37 double bonds per tail, and the *A.baumannii* membrane containing high levels of polyunsaturated fatty acids (up to 47%) and on average up to 1.02 double bonds per lipid tail). The differences in lipid composition corelate with differences in membrane properties such as the *E.coli* membrane being thicker and more ordered with slower lipid diffusion than the A.*baumannii* membrane.^19, 23^ The differences in properties between the *E.coli* and *A.baumannii* inner membranes further emphasizes the need to study bacterial membrane from a variety of species to understand the species specific differences in membrane properties. To create computational models of bacterial membranes, information about the membrane lipid composition is required. Such lipidomic data is available for the Gram-negative ESKAPE pathogens, *K.pneumoniae*, *P.aeruginosa*, E.*cloacae* and *E.coli*.^24,25^ These 4 species have variation in their phospholipid composition such that the concentration of CL ranges from 11% in *P.aeruginosa* to 3% in *E.cloacae*.^24, 25^ While the concentration of PG ranges from 21% in *P.aeruginosa* and *E.cloacae* to 5% in *K.pneumoniae*.^24, 25^ Finally, the PE concentration varies from 60% in *P.aeruginosa* to 82% in *K.pneumoniae*.^24, 25^ There are also variations in the lipid tails across the *K.pneumoniae*, *P.aeruginosa*, E*.cloacae* and *E.coli* inner membranes. Specifically, the number of unsaturations per lipid tail ranges ranging from 0.46 in *P.aeruginosa* to 0.18 in *K.pneumoniae*.^19, 26, 27^ Furthermore, *E.coli* and *P.aeruginosa* have fewer cyclopropane fatty acids than *K.pneumoniae or* E*.cloacae*.^19, 26, 27^ Based on the variation in lipid composition across Gram-negative bacteria, there is interest in studying the inner membrane of Gram-negative bacteria for a wide range of species in order to pinpoint how the different lipid compositions are impacting membrane properties.

The present study develops Martini coarse-grain models for the inner membrane of *K.pneumoniae*, *P.aeruginosa*, *E.cloacae* and *E.coli*. Through coarse-grain MD simulations, we elucidate the impact of varied lipid compositions on membrane properties across these Gram-negative bacterial species. Model membranes with simplified lipid composition were also studied to help pinpoint the effects of specific lipid species in modulating the properties of the membrane. Our findings highlight the critical role cardiolipin plays in modulating the structural properties of the Gram-negative bacteria inner membrane. The membrane models developed in this work contribute to a growing number of computational membrane models that replicate the composition of cell membranes and will facilitate future investigations into processes involving bacterial inner membranes, including the structure and function of transmembrane proteins.

## Methods

### Membrane composition

Four model bacteria were chosen for the current study namely *K.pneumoniae*, *P.aeruginosa*, *E.cloacae* and *E.coli*. Models of the inner membrane for each bacteria were constructed based on aggregated lipidomic data that provided information on both the lipid headgroup composition^24, 25^ and tail group composition^19, 26, 27^. The outer and inner leaflets were modeled as symmetric. Compositions of the membranes modeled in this work are provided in Figure 1 and Tables S1-S2. We note that although changes in membrane composition are known to occur based on factors such as diet and environment,^28–30^ these factors are not accounted for in the current work. Nevertheless, the current models provide insight into the effects of changing lipid composition on the biophysical properties of the inner membrane of Gram-negative bacteria.

**Figure 1.**
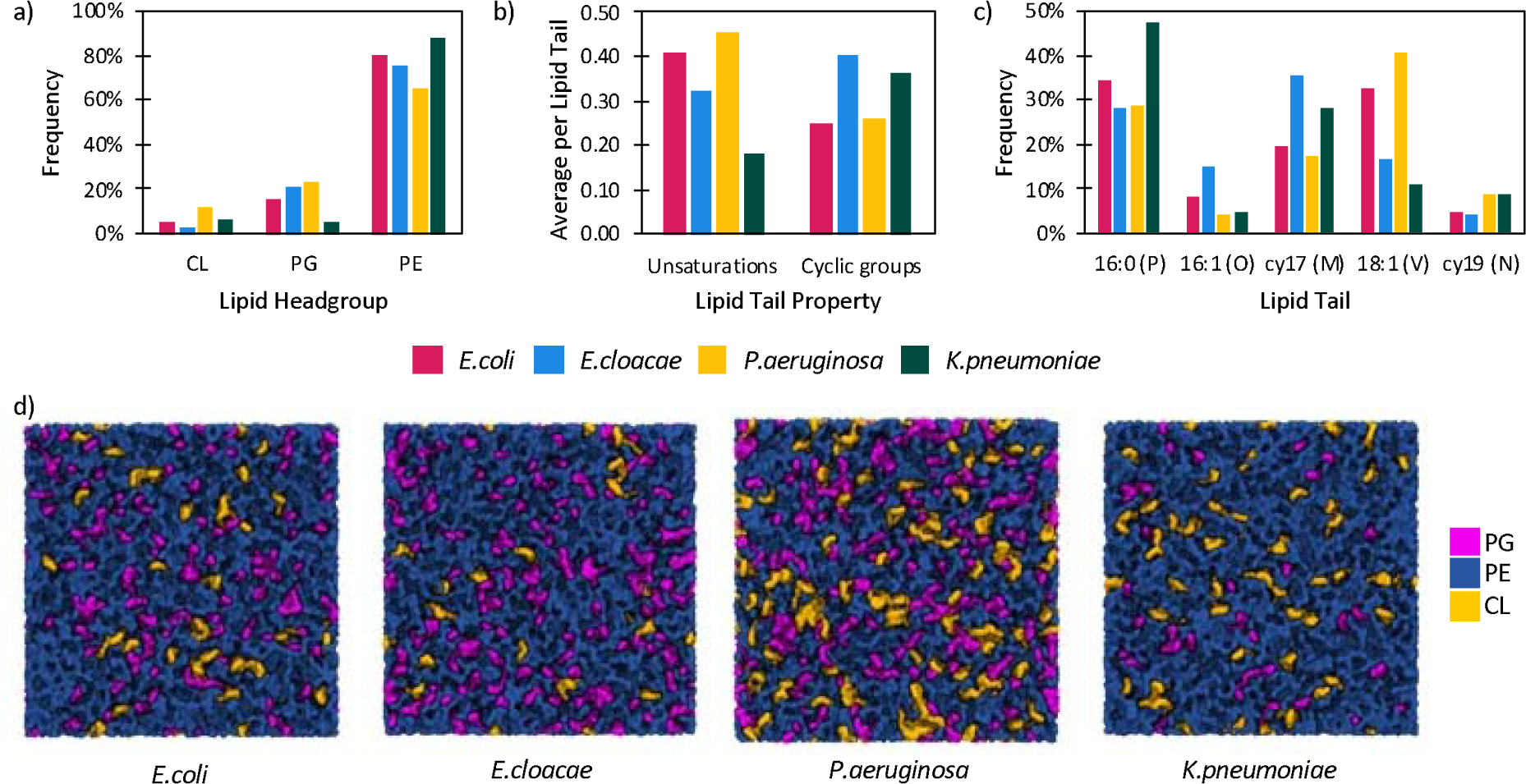
Composition of the *E.coli, E.cloacae*, *K.pneumoniae* and *P.aeruginosa* inner membranes. The composition has been displayed by a) lipid headgroup, b) lipid tail categories and c) fatty acid structure. d) A top-down view of the 4 bacterial membranes simulated in the current work colored by lipid class (PG in pink, PE in blue and CL in yellow).

Four model membranes were constructed that have variations in the lipid headgroup compositions but no variation in the lipid tail groups. Each lipid class was modeled by the single lipid species that was the most prevalent across the membranes studied. The resulting lipids have a mixture of saturated (P: 16:0), monounsaturated (V: 18:1), and cyclopropane (M: cy17) containing lipid tails. Since PE lipids are the most prevalent lipid class in the bacterial inner membrane (65-88%), PE was treated as the base lipid and the impact of PG and CL lipid concentration was investigated. The four model membranes are 1) high levels of PG (5% MPPV-CL, 25% PMPG, 70% PVPE), 2) low levels of PG (5% MPPV-CL, 5% PMPG, 90% PVPE), 3) high levels of CL (10% MPPV-CL, 20% PMPG, 70% PVPE) and 4) low levels of CL (5% MPPV-CL, 20% PMPG, 75% PVPE).

### Simulations

Martini 3.0 force field parameters^31^ were used in all simulation. Most lipid species were modeled based on previously published parameters,^19^ while additional PE and PG lipids parameters were created by manually combining head and tail group parameters of the previously published lipid species. All systems were 20 Å x 20 Å X 10 Å with the membrane aligned in the xy plane. Three replicates of each system were build using a custom version of the *insane* package.^32^ Each system contains ∼1340 lipids, ∼16 coarse grain waters per lipid, Na^+^ counter ions and 0.15 M NaCl. All simulations were performed using Gromacs 2023.1, using the recommended MARTINI simulation configuration, including the periodic boundary condition (PBC).^33^ The temperature (310 K) and pressure (1 bar) were maintained using the Bussi thermostat^34^ (τ_T_=1 ps) and semi-isotropic Parrinello-Rahman barostat^35, 36^ (τ_p_=12 ps and compressibility=3×10^-4^ bar^-1^), respectively. Each system underwent steepest descent minimization followed by 50 ns equilibration with an increasing timestep (10, 20 and 25 fs). Finally, production simulations of 10 µs were carried out using a 25 fs timestep.

### Analysis

LiPyphilic^37^ was used to analyze membrane thickness (distance between PO4 beads), area per lipid (APL), lipid order parameters, and lipid clustering. Membrane curvature was analyzed using MDAnalysis,^38, 39^ while Gromacs 2023.1 was used to analyse lipid diffusion. AmberTools23 was used to analyse water shells and VMD was used to analyze water permeation.

In analyzing water permeation, the water molecules were defined as above the membrane, below the membrane, in the membrane from above, or in the membrane from below. A permeation event occurred when a water molecule transitioned from “in the membrane from above” to “below the membrane” or “in the membrane from below” to “above the membrane”. Through including directionality when calculating membrane permeability any instances in which a water molecule bobs below the lipid headgroups and then back out into solution will not be considered a permeation event and any transitions across the PBC will also not be considered permeation events.

The isothermal membrane area compressibility modulus (K_A_) was calculated as:

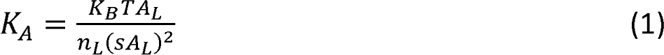

where K_B_ is the Boltzmann constant, T is the simulation temperature (310 K), A_L_ is the average APL for the membrane, n_L_ is the number of lipids in each leaflet, and sA_L_ is the standard deviation of the membrane APL.

## Results and Discussion

### Membrane thickness differs between the bacterial species, but the changes cannot be linked to one single factor of the membrane composition

The relationship between membrane thickness, lateral packing, and membrane fluidity determines the overall membrane dynamics. Changes in membrane thickness have been shown to have an impact on essential cellular processes, including small molecule permeability and membrane protein function.^40–43^ Our results indicate that variations in the composition of the bacterial inner membrane impact the membrane thickness. Specifically, the *E.coli* (40.2±0.2 Å) and *E.cloacae* (39.7±0.2 Å) inner membranes are significantly thicker than the *K.pneumoniae* (33.2±0.2 Å) and *P.aeruginosa* (32.7±0.2 Å) inner membranes (Figure 2a and S1). Previous simulations on a 14-component Avanti lipid *E.coli* inner membrane measured a thickness of 39 Å^19^ and experimentally, the *E.coli* membrane has been determined to 41 ± 1 Å thick at 298 K,^44^ which is within error of our measured thickness for the *E.coli* membrane (40.2±0.2 Å) and thereby justifies the accuracy of our computational methodology.

**Figure 2.**
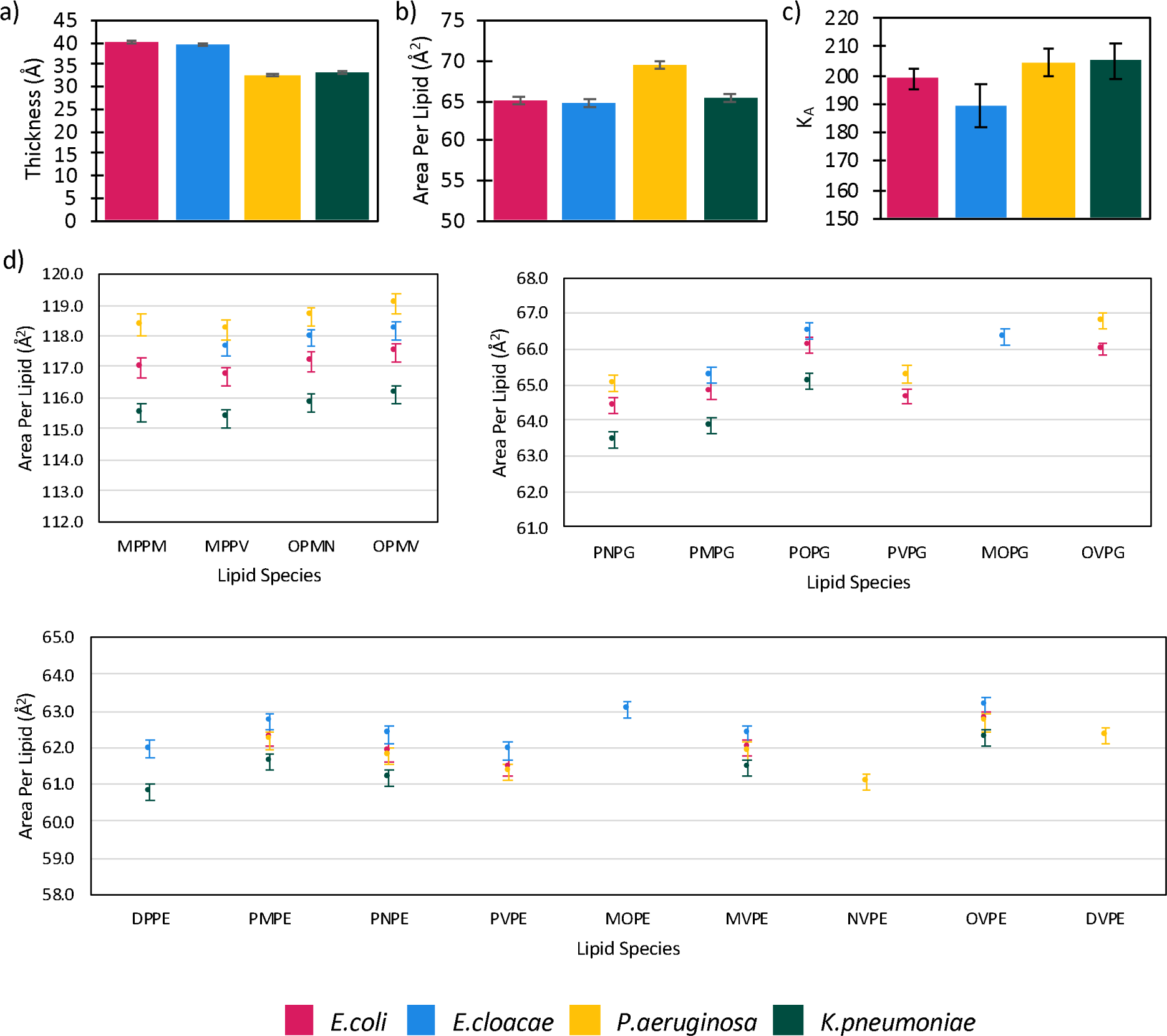
Overall membrane a) thickness, b) area per lipid and c) isothermal membrane area compressibility modulus (K_A_) of the *E.coli, E.cloacae*, *K.pneumoniae* and *P.aeruginosa* inner membrane derived from triplicate 10 μs coarse grain molecular dynamics simulations. d) The area per lipid for each individual lipid species during the simulations. Reported errors are the standard error of the mean across the three replicates.

Membrane thickness has been previously shown to be modulated by a variety of factors including phospholipid tail length, lipid tail order, and composition of the membrane.^45^ The differences in membrane thickness between *E.coli, E.cloacae*, *K.pneumoniae* and *P.aeruginosa* cannot be attributed to average lipid tail unsaturation (0.41, 0.32, 0.18 and 0.46 for *E.coli, E.cloacae*, *K.pneumoniae* and *P.aeruginosa*, respectively; Table S3) or length of the phospholipid tails (all membranes contain lipids with 16:0, 16:1, cy17, 18:1 or cy19 tails and all membranes have an average lipid tail length of 17 carbons). The concentration of CL in the membranes also does not explain the differences in membrane thickness, with the *P.aeruginosa* membrane having the most CL (12%) and the thinnest membrane, however the *E.coli* membrane is thicker than the *E.cloacae* membrane but has more CL (5% vs 3%). Overall, our data indicates that the compositional differences that occur within the inner membrane of bacteria can have significant impacts on the membrane thickness that are not directly linked to a single property of the membranes.

### Increased concentration of CL lipids leads to a larger area per lipid in the *P.aeruginosa* inner membrane than the other bacterial membranes studied

The *E.coli, E.cloacae*, and *K.pneumoniae* inner membranes have similar membrane fluidity with an overall area per lipid (APL) of 65.0 ± 0.5 Å^2^, 64.7 ± 0.5 Å^2^, and 65.2 ± 0.5 Å^2^, respectively. The *P.aeruginosa* inner membrane displays greater fluidity with an overall APL of 69.4 ± 0.5 Å^2^ (Figure 2b and S1). Using the measured APL, the isothermal membrane area compressibility modulus (K_A_) can be measured. A greater K_A_ is seen for the *K.pneumoniae* and *P.aeruginosa* inner membranes (204.5±5.1 and 205.0±6.4 mN m^-1^, respectively) than the *E.coli* and *E.cloacae* inner membranes (198.8±3.7 and 189.4±7.4 mN m^-1^, respectively; Figure 2c). The higher rigidity of the *K.pneumoniae* and *P.aeruginosa* inner membranes correlates with the thinner membranes observed for these systems.

When the APL is considered by individual lipid species, significant differences are observed between the lipid species (Figure 2d). CL lipids have the largest APL (115-119 Å^2^), while PG lipids have a larger APL than PE (63-67 Å^2^ and 61-63 Å^2^, respectively). While not directly comparable due to being measured in single lipid species membranes our simulations agree with the experimental trends in lipid APL. Specifically, experimentally POPE has an APL of 58 Å^2^ (at 308 K),^46^ POPG has an APL of 61 Å^2^ (at 303 K)^47^ and tetraoleoylcardiolipin has an APL of 130 Å^2^ (at 303 K).^48^ The APL of CL lipids is less than double the APL observed for phospholipid species, therefore per phospholipid headgroup, the APL of CL is smaller than both PG and PE. For all lipid classes, *K.pneumoniae* displays the lowest APL, followed by *E.coli*. *P.aeruginosa* has the largest APL for CL, while *E.cloacae* has the largest APL for PG and PE lipids. Differences in membrane composition influence the overall APL in the membrane. Specifically, the high CL composition of the *P.aeruginosa* membrane coupled with the higher APL for CL species leads *P.aeruginosa* to have the highest APL.

Further differences are observed when the lipid tails are considered (Figure 2d). As expected for lipids without cyclopropane groups in their tails, increasing levels of unsaturation increase the APL.^49, 50^ For example, across all membranes DPPE with two saturated lipid tails, has a lower APL than OVPE, which has two monounsaturated lipid tails, and POPG with a single monounsaturated lipid tail has a lower APL than OVPG, which has two monounsaturated lipid tails. Clear trends in APL relative to lipid tail unsaturation are not always observed for lipids containing cyclopropane groups. Previous research has indicated that cyclopropane rings lead to an increase in APL compared to saturated lipids^21, 51^ and the effect of a single cyclopropane ring is similar to that of two double bonds for PC lipids.^51^ In the current study we show that for CL lipid species a cyclopropane group often appears to have a similar effect to double bond in terms of effects on APL. Specifically, OPMV-CL with 1 cyclopropane fatty acid tail and 2 monounsaturated fatty acid tails has the largest APL in all membranes, while MPPV-CL with 1 cyclopropane fatty acid tail and 1 monounsaturated fatty acid tails and MPPM-CL with 2 cyclopropane fatty acid tail have a similar APL that is smaller than OPMV-CL in all membranes. Conversely for PG lipid species, lipids with a single cyclopropane fatty acid tail (PNPG and PMPG) have a smaller APL than lipids with a single monounsaturated fatty acid tail (PVPG and POPG). Therefore, there is a lipid class dependency for the impact of the cyclopropane group on the resulting APL.

### Lipid order changes in response to variations in lipid tail unsaturation

Lipid order parameter is a biophysical property originally derived from NMR, which measures the order of the membrane through calculating the conformations the lipid tails adopt.^52^ A lipid order of 1 indicates perfect alignment of the lipid tails with the membrane normal, while a value of –0.5 indicates that the lipid tails are perpendicular to the membrane normal and a value of 0 indicates a random distribution.^37^ The *K.pneumoniae* membrane displays the highest order (0.38 ± 0.01), while *E.cloacae* (0.34 ± 0.01), *E.coli* (0.35 ± 0.01), and *P.aeruginosa* (0.35 ± 0.01) are all less ordered (Figure 3a and S2). Membrane order is known to be influenced by lipid tail unsaturation, with higher levels of unsaturation leading to a decrease in membrane order.^49, 50^ The observed trend in membrane order is consistent with the trend in lipid tail unsaturation for the 4 bacterial membranes. Specifically, *K.pneumoniae* membrane has 0.18 unsaturations per lipid tail, while the *E.cloacae*, *E.coli*, and *P.aeruginosa* membranes contain 0.32, 0.41 and 0.46 unsaturations per lipid tail, respectively. All membranes modeled are more ordered than what was previously observed for the inner membrane of *A.baumannii* (0.26-0.29 Å), which contains higher levels of lipid unsaturation (0.71-1.02 unsaturations per lipid tail).^23^

**Figure 3.**
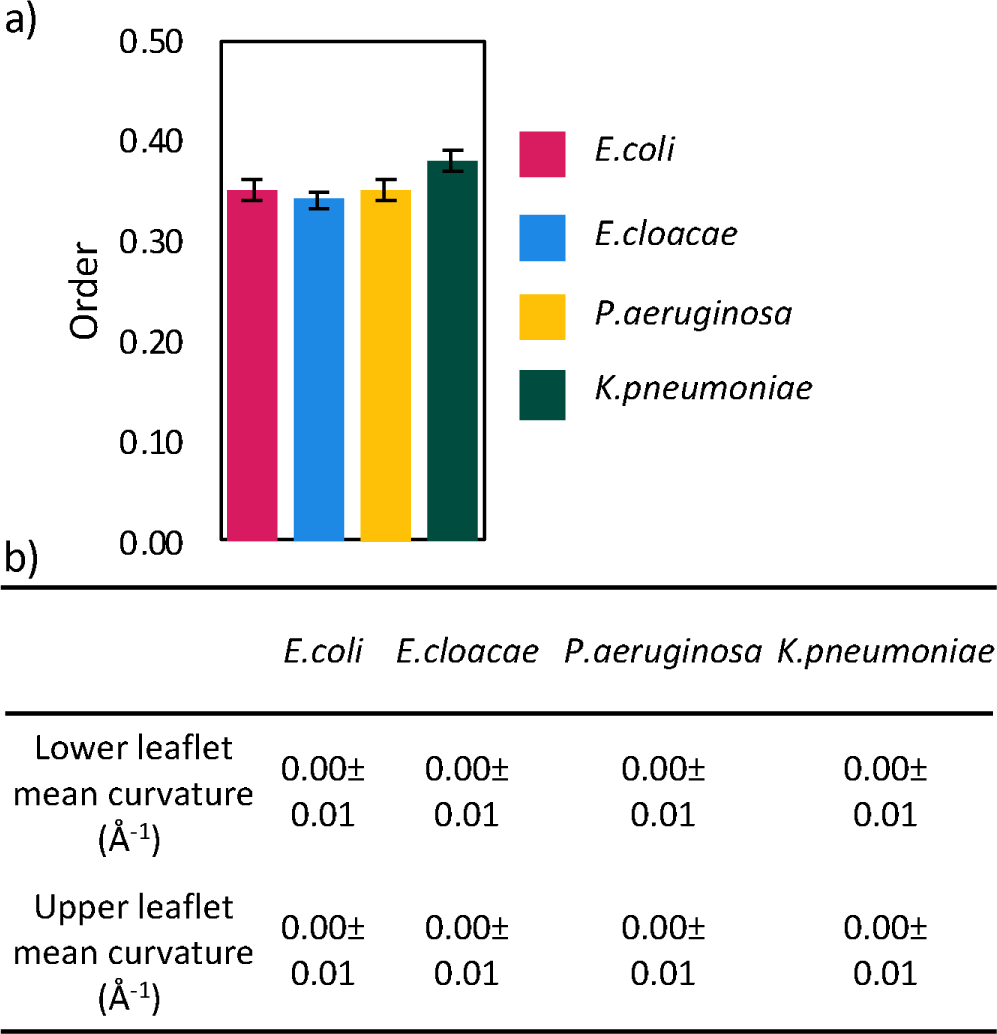
Average a) lipid order and b) membrane curvature in the lower and upper leaflets of the *E.coli, E.cloacae*, *K.pneumoniae* and *P.aeruginosa* inner membrane over the course of triplicate 30 μs of coarse grain molecular dynamics simulation. Reported errors are the standard error of the mean across the three replicates.

When membrane order is considered for the individual lipid species, the more saturated lipids generally have a higher membrane order (Figure S3). The saturated lipid DPPE consistently shows the highest order (0.42-0.45), while MOPE and MOPG with one site of unsaturation and one cyclopropane group show the lowest membrane order (0.29). Phospholipids with 2 sites of unsaturation (OVPE, DVPE, and OVPG) show similar order (0.30-0.31) to MOPE and MOPG. OPMV-CL, which contains 1 cyclopropane group and 2 sites of unsaturation, has a slightly higher order (0.33-0.35) than the phospholipids with 2 sites of unsaturation. Across the 4 membranes, each lipid species shows a consistent average order (maximum range of 0.03). Despite the small range, the order for lipid species in the *K.pneumoniae* membrane is generally the highest, followed by *E.coli* and *P.aeruginosa* and lowest in *E.cloacae*. This indicates that the interactions taking place between the lipids within each membrane are further modulating the lipid order beyond the effects simply observed due to the chemical structure of the lipid.

### Membrane curvature is not impacted by lipid composition

Membrane lipid composition is known to effect membrane curvature.^53^ Specifically, the interplay between lipid tail volume and headgroup volume influences whether a lipid is likely to induce a negative or positive curvature. PE is considered a conical lipid, prone to inducing negative curvature due to the small lipid headgroup and typical unsaturated nature of the lipid tails.^54^ PG is a lamellar lipid meaning that PG has a similar sized headgroup and tail group volume and this does not induce curvature.^54^ Finally, cardiolipin is conical lipid that favours negative curvature and has been shown to localizes in areas to induce curvature.^55^ Therefore, based on the predominance of PE and CL lipids it would be expected to see negative curvature in the membranes. Despite the differences in membrane composition, there is no difference observed in the membrane curvature across the four bacteria membranes (Figure 3b and S4). Similar to observed in previous work on the *E.coli* inner membrane,^19^ all modeled membranes adopt a planar geometry (mean membrane curvature of 0.00±0.01 Å^-1^ for both the upper and lower leaflets). Despite the difference in membrane headgroup composition, the lipid tail group composition is less varied between the membranes, with all membrane containing on average less than 0.5 site of unsaturation or cyclopropane groups per lipid. The saturated nature of the modeled bilayers may explain the lack of membrane curvature. Notably, previous work on the PUFA enriched *A.baumannii* inner membrane that contain up to 1 site of unsaturation per lipid contain a 9-10° deviation from the bilayer normal.^23^

### Lipid diffusion depends on lipid class and membrane composition

Changes in lipid diffusion have been linked with changes in membrane curvature, lipid order and lipid domain formation.^43, 56, 57^ Although we saw no variation in curvature and only slight differences in lipid order, the lipid lateral self-diffusivity observed in the current work varies across the bacterial species. The *E.cloacae* inner membrane shows the highest rate of lipid lateral self-diffusivity (5.93 ± 0.44 x 10^-7^ cm^2^/s), followed by the *E.coli* (5.76 ± 0.17 x 10^-7^ cm^2^/s), *K.pneumonia* (5.73 ± 0.33 x 10^-7^ cm^2^/s) and *P.aeruginosa* (5.37 ± 0.25 x 10^-7^ cm^2^/s) inner membranes (Figure 4a). The trend in overall lipid lateral self-diffusivity correlates with the concentration of CL lipids present in the membrane with the *E.cloacae* membrane having the lowest concentration (3% CL), followed by *E.coli* (5% CL), *K.pneumonia* (7% CL) and *P.aeruginosa* (12% CL) membranes. We do not see any impact of overall lipid tail unsaturation levels on lipid lateral self-diffusivity. When the lipid lateral self-diffusivities are calculated by lipid class (Figure 4b), across all membranes the phospholipid species display higher rates of lateral self-diffusivity (5.4-6.0 x 10^-7^ cm^2^/s) than CL species (4.0-4.8 x 10^-7^ cm^2^/s), which at least in part explains why membranes with higher levels of CL show lower overall lipid lateral self-diffusivity. The *E.cloacae* membrane shows the highest rate of lipid lateral self-diffusivity for each lipid class, while the *P.aeruginosa* membrane shows the lowest rate of lipid lateral self-diffusivity for each lipid class. For example CL in the *E.cloacae* membrane display a lipid lateral self-diffusivity of 4.81 ± 0.99 x 10^-7^ cm^2^/s, while CL in the *P.aeruginosa* membrane show a lipid lateral self-diffusivity of 4.07 ± 0.33 x 10^-7^ cm^2^/s. The variations seen within a lipid class across the membranes, points to compositional effects of lipid diffusion that may be likely linked to lipid domain formation.

**Figure 4.**
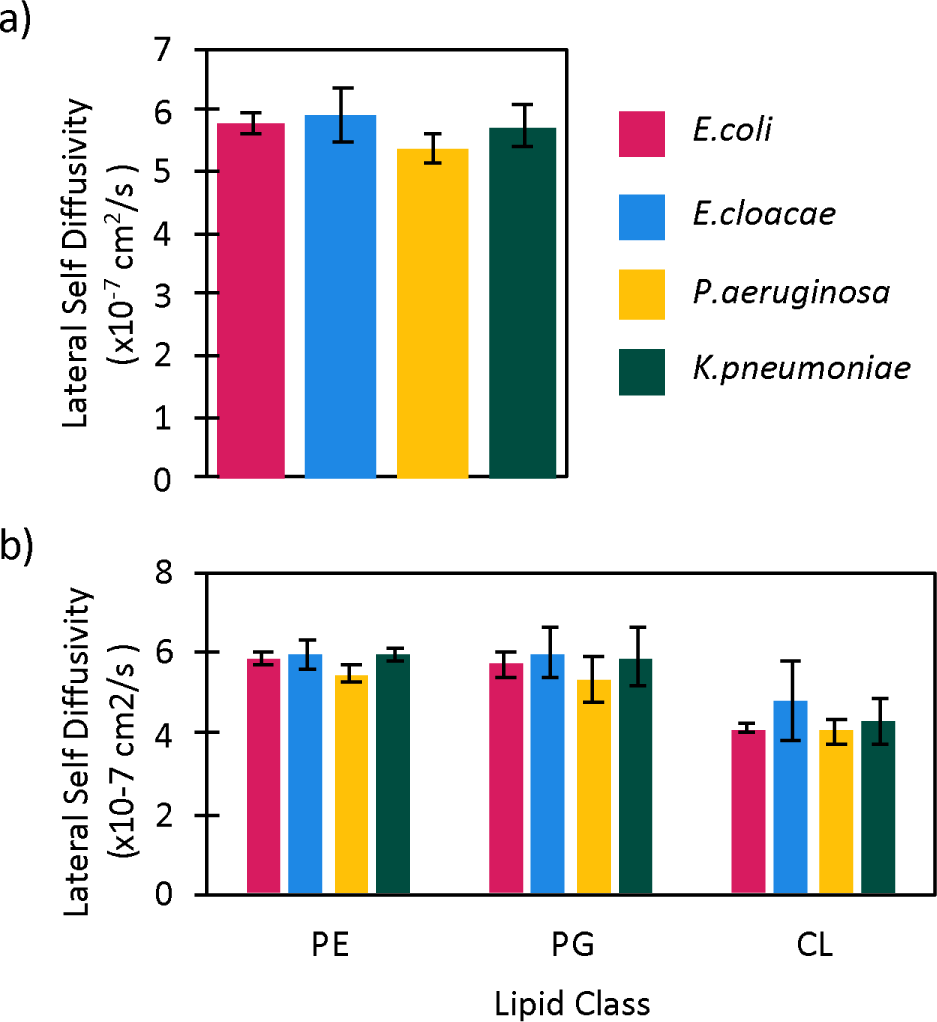
Lipid lateral self-diffusivity averaged over a) all lipid species or b) by lipid headgroup class for the *E.coli, E.cloacae*, *K.pneumoniae* and *P.aeruginosa* inner membranes over the course of triplicate 30 μs of coarse grain molecular dynamics simulations. Reported errors are the standard error of the mean across the three replicates.

### Lipid domain formation is impacted by the CL concentration of the bacterial inner membrane

Changes in membrane composition have been shown to affect the formation of lipid raft domains.^23, 58–60^ To understand how changes to the membrane compassion across bacterial species effects the formation of lipid raft domains, the lipid enrichment-depletion indexes were calculated for the 4 bacterial species. A lipid enrichment-depletion index represents the propensity for two lipid species to colocalize within the membrane.^61^ If the lipid enrichment-depletion index of two lipid species is 1, it means the two lipid species are randomly distributed throughout the membrane, while if the index is >1 the two lipid species are co-localized or enriched with respect to one another, while if the index is < 1 the two lipid species are avoiding one another or depleted with respect to one another. Previous research has indicated that lipids tend to localize based on electrostatic interactions between lipid headgroups.^58, 62^ In the current work we observe clustering between zwitterionic PE lipids and di-anionic CL lipids (enrichment-depletion index= 1.3 to 1.4), with the greatest enrichment observed for *E.cloacae* and the lowest levels of enrichment in *P.aeruginosa* (Figure 5a). Like other membrane properties the differences in PE:CL clustering aligns with the concentration of CL in the bilayers, with higher levels of CL correlating with less PE:CL clustering. Furthermore, all other lipid contacts are neither enriched nor depleted with contact fractions between 0.8 and 1.2. The formation of CL:PE lipid domains and the lack of domain formation with other lipid classes has been previously reported in *E.coli*^19^ and *A.baumannii*^23^ inner membranes. Beyond clustering based on lipid headgroups, previous work on the impact of polyunsaturated fatty acid on the properties of the *A.baumannii* inner membrane indicated that lipid tail saturation can also impact lipid enrichment-depletion indexes.^23^ In the current work we see no trends in lipid clustering based on the properties of the lipid tails, with all average lipid enrichment-depletion index between 1.0-1.1 when averaged based on lipid tail saturation (Figure 5b). The lack of lipid clustering observed in the current work is likely due to the low levels of lipid unsaturation.

**Figure 5.**
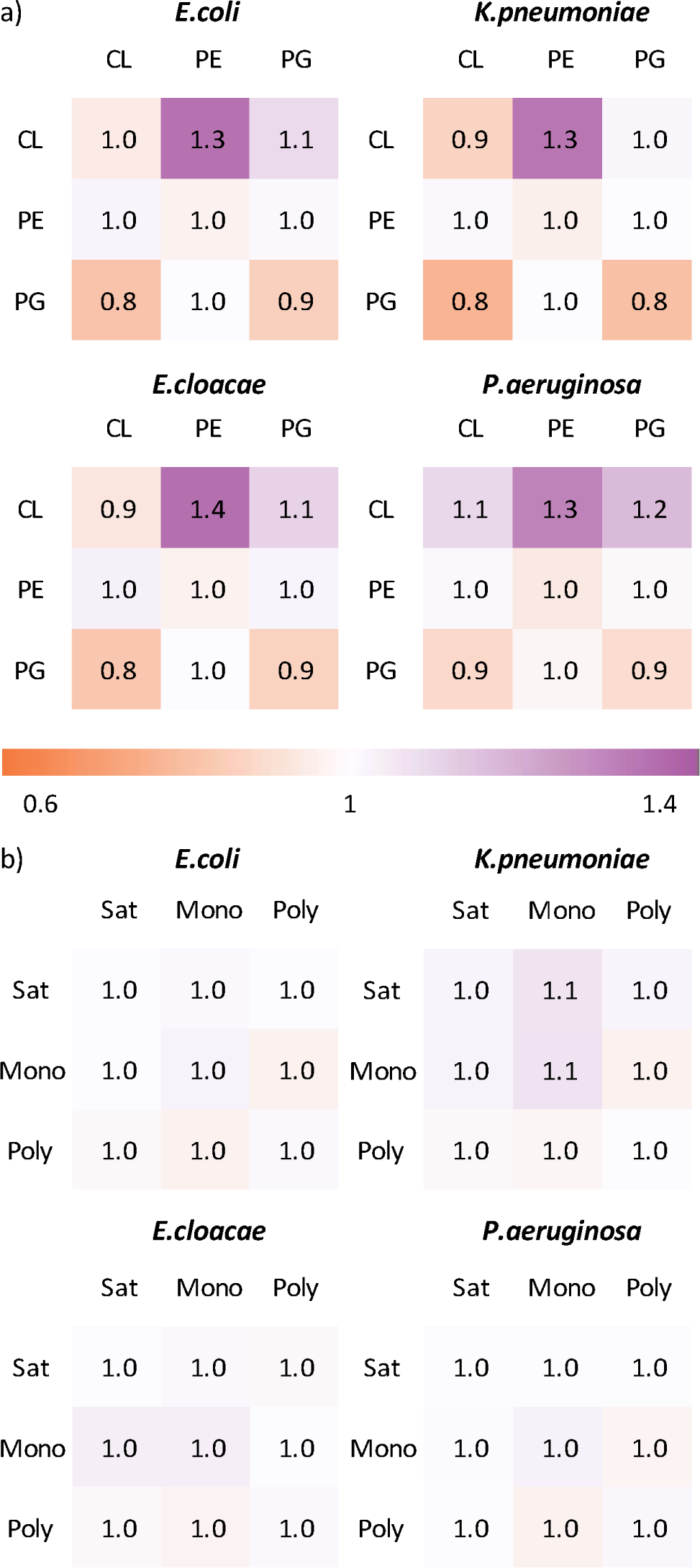
Depletion enrichment index (DEI) reporting lipid clustering in the *E.coli, E.cloacae*, *K.pneumoniae* and *P.aeruginosa* inner membranes over the course of triplicate 30 μs of coarse grain molecular dynamics simulation. The values have been averaged based on lipid a) headgroup or b) tail. All reported DEI values have a standard error of the mean of less than 0.02.

When the lipid clustering is considered by individual lipid species, the same general trends are maintained that are observed for the lipid classes (Figure S5-S8). For *P.aeruginosa,* all DEI are within 0.1 of the average values for the lipid class. Interestingly, however for *E.coli, E.cloacae* and *K.pneumoniae* the self-interactions between CL:CL are depleted, while the interactions between differing CL species are neither enriched nor depleted. Compared to *P.aeruginosa*, the *E.coli, E.cloacae* and *K.pneumoniae* bilayers all have lower levels of CL (3-6% vs 12%) and therefore the observed self-depletion of CL:CL interactions may allow for the CL to interact with other lipid species such as the observed clustering with PE. Furthermore, for *E.coli* and *K.pneumoniae* the same depletion of self-interactions is observed for PG lipids. *E.coli* and *K.pneumoniae* have the lowest concertation of PG lipids, and therefore the observed depletion of self-interactions may occur to allow the PG lipids to interact with other lipid species.

### *P.aeruginosa* has greater water penetration than the other bacterial membranes studied

One of the purposes of a lipid bilayer is to create a hydrophobic barrier around a cell or organelle in order to create an environment that is conducive for life. Although this barrier serves to prevent the movement of material into and out of the cell, water and other small molecules can permeate the membrane without disrupting the overall structure.^63^ In order to understand how water is associating with the 4 bacterial inner membranes studied, we calculated the number of water molecules associated with the lipid headgroups as well as the number of water molecules that permeate the bilayer. More water is associated with and penetrates the *P.aeruginosa* bilayer than the *E.coli, E.cloacae* and *K.pneumoniae* bilayers (Figure 6). Specifically, the *E.coli, E.cloacae* and *K.pneumoniae* bilayers have on average 0.86 to 0.87 ± 0.02 coarse grain water molecules within 5 Å of each lipid headgroup, while the *P.aeruginosa* bilayer has on average 0.92 ± 0.02 coarse grain water molecules within 5 Å of each lipid headgroup. Similarity, while the *E.coli, E.cloacae* and *K.pneumoniae* bilayers have on average 4.14±0.04 coarse grain water molecules within 8 Å of each lipid headgroup, while the *P.aeruginosa* bilayer has on average 4.43 ± 0.04 coarse grain water molecules within 8 Å of each lipid headgroup. Greater water penetration is also observed for the *P.aeruginosa* bilayer compared to the *E.coli, E.cloacae* and *K.pneumoniae* bilayers, with < 1.5 permeation events observed per water molecule over the course of the 10 μs simulation for the *E.coli, E.cloacae* and *K.pneumoniae* bilayers, but 3.4 permeation events are observed per water molecule over the course of the 10 μs simulation for the *P.aeruginosa* bilayer. Notably, water molecules penetrate lipid bilayers without disrupting the structure of the membrane through occupying holes in the lipid matrix. The *P.aeruginosa* bilayer has greatest APL, and therefore has the most space for water molecules to occupy without disrupting the bilayer.

**Figure 6.**
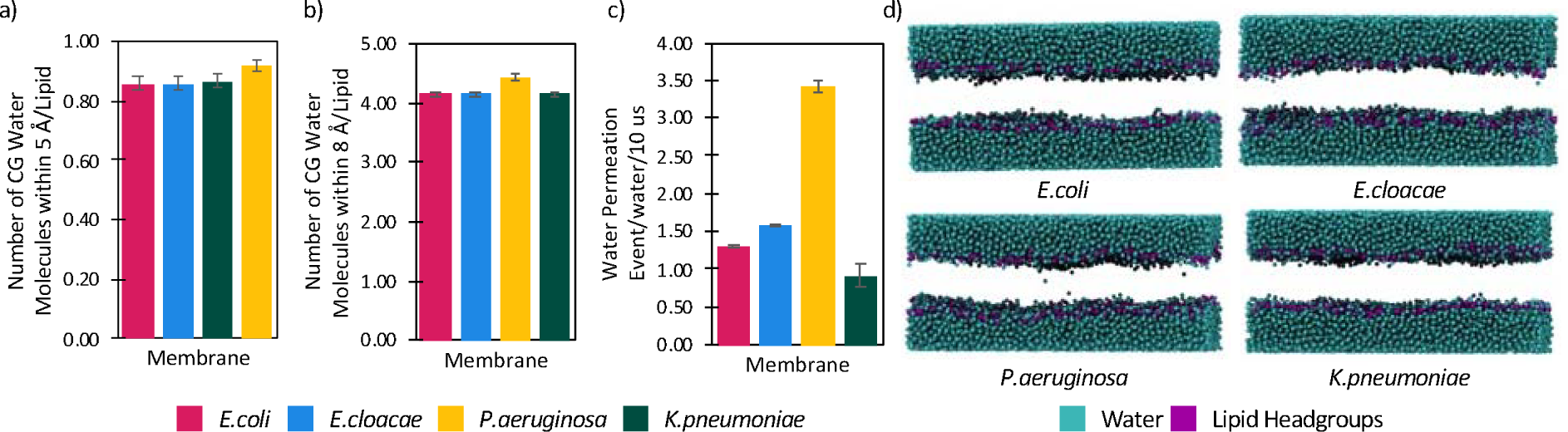
Water associates more with the *P.aeruginosa* membrane over the course of the molecular dynamics simulations than the other membranes studied as monitored by number of water molecules within a) 5 Å or b) 8 Å of the lipid headgroups, as well as c) the number of coarse grain water molecules that penetrate the membrane over the course of the 10 μs simulation. d) Snapshots showing the number of water molecules in the center of the bilayer at the end of a single replicate of a 10 μs simulation.

### Model membranes without the full complexity of the lipid tails are not able to reproduce the properties of the complex models of the inner membrane but highlight the role of CL in dictating membrane properties

Throughout our analysis of the bacterial inner membranes, many properties such as APL, lipid clustering, lipid diffusion, and water penetration were shown to be impacted by lipid headgroup composition. To try and further pinpoint the origin of the variations observed in the membranes, model membranes were constructed that had variations in the lipid headgroup compositions but no variation in the lipid tail groups. Since PE lipids make up most of the membrane composition in the bacterial inner membrane (65-88%), PE was treated as the base lipid and the impact of PG and CL lipid composition was investigated. Membranes with high levels of PG (5% MPPV-CL, 25% PMPG, 70% PVPE) mimic the composition of the *E.cloacae* and *E.coli* membranes, while low levels of PG (5% MPPV-CL, 5% PMPG, 90% PVPE) mimic the *K.pneumoniae* membrane. Furthermore, a membrane with high levels of CL (10% MPPV-CL, 20% PMPG, 70% PVPE) mimic the composition of the *P.aerugonosa* membrane, while a membrane with low levels of CL (5% MPPV-CL, 20% PMPG, 75% PVPE) mimics the composition of the *E.cloacae* and *E.coli* membranes.

The changes in the membrane thickness and lipid order that were observed in the 4 bacterial membranes were not attributed to changes in the lipid headgroup composition. Therefore, it is not surprising that the membrane thickness did not change upon increasing the levels of CL lipid concentration (40.9±0.2 Å vs 41.0±0.2 Å) or PG lipid concentration (41.1±0.2 vs 40.9±0.2 Å, Table 1). All model membranes display thicknesses similar to the *E.cloacae* or *E.coli* membranes, which are significantly thicker than the *K.pneumoniae* and *P.aerugonosa* membranes. Further indicating that lipid tail composition plays a major role in determining membrane thickness and that the lipid tail group diversity needs to be fully represented to reproduce the thickness of the bacterial inner membranes. Similarly, no change was observed in lipid ordering upon modifying the composition of the model membranes with all membranes displaying a lipid order of 0.39. The observed lipid order for the model membranes is slightly greater than observed for the complex membrane systems (by 0.01-0.05). As lipid order measures the packing of the lipid tails it makes sense that this property would be affected by changes to the lipid tail composition.

**Table 1.** Calculated membrane properties from triplicate 10 μs coarse grain molecular dynamics simulations of model bacterial membranes with varying concertation of PG or CL lipids.

|  | *Low-CL* | *High-CL* | *Low-PG* | *High-PG* |
| --- | --- | --- | --- | --- |
| Composition | 5% MPPV-CL, 20% PMPG, 75% PVPE | 10% MPPV-CL, 20% PMPG, 70% PVPE | 5% MPPV-CL, 5% PMPG, 90% PVPE | 5% MPPV-CL, 25% PMPG, 70% PVPE |
| Unsaturation per lipid tail | 0.4 | 0.4 | 0.5 | 0.4 |
| Thickness ($\text{\AA}$) | $40.9 \pm 0.2$ | $41.0 \pm 0.2$ | $41.1 \pm 0.2$ | $40.9 \pm 0.2$ |
| Lipid order | $0.39 \pm 0.01$ | $0.39 \pm 0.01$ | $0.39 \pm 0.01$ | $0.38 \pm 0.01$ |
| Area per lipid ($\text{\AA}^2$) | $63.7 \pm 0.4$ | $66.7 \pm 0.4$ | $63.4 \pm 0.4$ | $63.9 \pm 0.4$ |
| Lipid lateral self-diffusivity ($\times 10^{-7} \text{ cm}^2/\text{s}$) | $5.37 \pm 0.32$ | $5.22 \pm 0.38$ | $5.73 \pm 0.18$ | $5.52 \pm 0.22$ |
| Lower leaflet mean curvature ($\text{\AA}^{-1}$) | $0.00 \pm 0.01$ | $0.00 \pm 0.01$ | $0.00 \pm 0.01$ | $0.00 \pm 0.01$ |
| Upper leaflet mean curvature ($\text{\AA}^{-1}$) | $0.00 \pm 0.01$ | $0.00 \pm 0.01$ | $0.00 \pm 0.01$ | $0.00 \pm 0.01$ |
| Number of water molecules within 5 $\text{\AA}$ of the lipid headgroups (per lipid) | $0.85 \pm 0.02$ | $0.89 \pm 0.02$ | $0.84 \pm 0.02$ | $0.86 \pm 0.02$ |
| Number of water molecules within 8 $\text{\AA}$ of the lipid headgroups (per lipid) | $4.06 \pm 0.04$ | $4.25 \pm 0.04$ | $4.05 \pm 0.04$ | $4.07 \pm 0.04$ |

In the model membranes little variation is observed in the lipid domain formation and membrane curvature. Specifically, for the high and low CL membranes PE:CL domains are enriched with a lipid contact fraction of 1.3, while the remainder of the lipid contact fractions are 1.0±0.1. In the low and high PG model membranes, PE:CL domains are enriched with a lipid contact fraction of 1.3, while the PG:PG contacts are depleted with low PG concentration (0.8 compared to 0.9 in the high PG membrane) and the CL:PG contacts are enriched with high PG concentration (1.2 compared to 1.0 in the low PG membrane). The remainder of the lipid contact fractions in the PG model membranes are 1.0±0.1. The clustering of zwitterionic PE lipids and di-anionic CL lipids aligns with the domain formation observed in the bacterial membrane systems and previous literature.^19, 23^ Additionally, as seen in the bacterial model membranes, little change is observed in membrane curvature. Specifically, all model membranes were planar with a mean membrane curvature of 0.00±0.01 Å^-1^. Therefore, while lipid head group plays a role within determining the curvature and domain formation within a membrane, the model lipid compositions studied in the current work have little impact either property.

Increased levels of cardiolipin were predicted to accounts for the larger APL, greater water penetration and lower lipid lateral self-diffusivity observed for the *P.aeruginosa* inner membrane compare to the other bacterial membranes studied. Consistent with this prediction, the high concentration cardiolipin model membrane has a larger APL (65.0±0.4 Å^2^, Table 1) than the low cardiolipin concentration model membrane (63.7±0.4 Å^2^). Furthermore, changing the concentration of PG in the model membranes has little impact on the overall APL of the membranes (63.4±0.4 vs 63.9±0.4 Å^2^ for the low and high concentration PG model membranes, respectively). Nevertheless, the complex bacterial membranes typically had a larger APL (64.7 to 69.4 Å^2^) than observed in the model membranes indicating that along with cardiolipin impacting the APL of the membranes, the levels of unsaturation in the lipid tail groups are further modulating the APL. In the model membranes the high CL membrane has a greater number of water molecules associated with the membrane (0.89±0.02 and 4.25±0.04 CG water molecules per lipid within 5 and 8 Å of each lipid headgroup, respectively, Table 1) than the low CL membrane (0.85±0.02 and 4.06±0.04 CG water molecules per lipid within 5 and 8 Å of the lipid headgroup, respectively). Consistent with the prediction that CL concentration is modulating the water surrounding the membrane and resulting permeation events, changing the concertation of PG lipids did not affect the number of water molecules associated with the lipids (0.84 to 0.86±0.02 and 4.05 to 4.07 ±0.04 CG water molecules per lipid within 5 and 8 Å of each lipid headgroup, respectively). Finally, based on the lipid lateral self-diffusivity, we proposed that increasing levels of CL leads to lower overall lipid lateral self-diffusivity. In the model membranes we observe lower lipid lateral self-diffusivity upon increased levels of CL (5.37 x 10^-7^ cm^2^/s and 5.22 x 10^-7^ cm^2^/s for the high and low CL membranes, respectively, Table 1). Notably however, we also observe lower levels of overall lipid lateral self-diffusivity upon increasing the levels of PG in the membranes (5.52 x 10^-7^ cm^2^/s and 5.73 x 10^-7^ cm^2^/s for the high and low PG membranes, respectively). Indicating that not only CL but also PG lipid concentration impacts the lipid lateral self-diffusivity. Furthermore, none of the model membranes can reproduce the overall lipid lateral self-diffusivity seen in the bacterial inner membrane.

## Conclusion

In the current study we have used available lipidomics data to generate computational models of the inner membrane of 4 ESKAPE bacterial species, namely *E.cloacae*, *E.coli*, *K.pneumoniae* and *P.aerugonosa*. While previous work had highlighted the differences in lipid composition across these pathogenic bacterial species, the impact of the compositional differences on membrane biophysical properties had not previously been explored. Model membranes that vary in the composition of the lipid classes (CL, PE and PG) were also investigated to help pinpoint the origin of differences in membrane properties observed across the 4 bacterial species.

*E.coli* and *E.cloacae* are listed as priority 1 or critical for the development of new antibiotics by the WHO. Both bacterial have similar biophysical properties of the inner membrane such as a membrane thickness of ∼40 Å, an area per lipid of ∼65 Å^2^, and low water permeability. The similarities in membrane properties observed in *E.coli* and *E.cloacae* is likely due to the similarity in membrane composition, including low CL concentration (<5%) and 0.3-0.4 unsaturations per lipid tail. Nevertheless, while addition of cyclopropane fatty acids to membrane models^20, 21^ enhances membrane fluidity, increasing the concentration of cyclopropane fatty acids from 0.25/lipid in the *E.coli* inner membrane to 0.40/lipid in the *E.cloacae* inner membranes does not appear to impact membrane properties. Conversely, the *P.aeruginosa*, which is listed as priority 2 by the WHO for the development of new antibiotics has an inner membrane with higher levels of cardiolipin than the *E.coli* and *E.cloacae* membranes but similar levels unsaturation and cyclopropane groups to the *E.coli* inner membrane. The difference in membrane composition of the *P.aeruginosa* inner membrane lead to a thinner membrane with a higher area per lipid and greater water permeability than the *E.coli* and *E.cloacae* inner membranes. Nevertheless, cardiolipin concentration alone is not able to explain the differences observed in the bacterial inner membranes. Specifically, the *K.pneumoniae* inner membrane has similar membrane thickness to the *P.aeruginosa* inner membrane yet has less cardiolipin (7% vs 12%, respectively). Regardless of the concentration of cardiolipin in the 4 bacterial membranes investigated, clustering is observed between cardiolipin and PE lipids, similar membrane order and almost no membrane curvature is observed. The lack of variation in lipid order, lipid domain formation and curvature is likely in part due to the low levels of unsaturation in the *E.cloacae*, *E.coli*, *K.pneumoniae* and *P.aerugonosa* bacterial inner membranes. Indeed, the *A.baumanni* which has higher levels of unsaturation but no cyclopropane lipids, is less ordered, more curved and shows variation in lipid domain formation with respect to levels of lipid tail unsaturation.^23^

The model membranes provide further evidence that APL, greater water penetration, lipid domain formation and lower lipid lateral self-diffusivity are influenced by CL concentration, while membrane thickness, order, and curvature are not directly impacted by CL concentration. Furthermore, the model membranes show that cardiolipin concentration alone is not able to account for all the observed differences in membrane properties. Additionally, like noted in previous studies on the *E.coli*,^19^ average plasma,^60^ neuronal^59, 60^ and epithelial^64^ membranes, lipid diversity is critical in order to accurately represent the biophysical properties of a membrane. In the case of the bacterial membranes representing the diversity of the lipid headgroups is not enough to capture the biophysical properties of the membranes. Therefore, future work on biological membrane systems should strive to represent the membrane with as much detail as possible.

Overall, this study underscores the significance of membrane composition in defining the properties of the bacterial inner membrane. Specifically, the concentration of cardiolipin plays a key role in determining the membrane fluidity and water permeability of the bacterial inner membrane. As cardiolipin concertation is related to key membrane properties it may also be impacting the ability for AMP to disrupt bacterial membranes. The models developed in this research can be utilized in future investigations to better understand the proteins embedded in the bacterial inner membrane. Additionally, this work lays the groundwork for innovative approaches to disrupting the bacterial membrane, offering potential new treatments for antibiotic-resistant infections.

## Data Availability Statement

The data that support the findings of this study are openly available at github.com/WilsonLabMUN/bacterial-inner-membrane.

## Supporting Information

Table S1: phospholipid species in the bacterial membrane models, Table S2-S3: head group and tail group composition of the bacterial membrane models, Figures S1: membrane thickness and APL over time, Figure S2: order by lipid species, Figure S3: order over time, Figure S4: membrane curvature mapped to membrane surface, Figure S5-S8: lipid enrichment-depletion index by individual lipid species.

## Acknowledgements

This work was funded by the Natural Sciences and Engineering Research Council (NSERC) of Canada (2023-05903) and Memorial University of Newfoundland. We would like to thank Nisarg Dave for their preliminary calculations related to this work. The research was undertaken with the assistance of resources and services from the Digital Research Alliance of Canada.

The authors declare no competing interests.

## Author Contributions

K.A.W. designed the research and wrote the paper; G.G., L.P., and S.R. performed the simulations; G.G., L.P., S.R. and K.A.W. contributed to the analysis. All authors have given their final approval to this version of the manuscript.

## Table of Contents Graphic

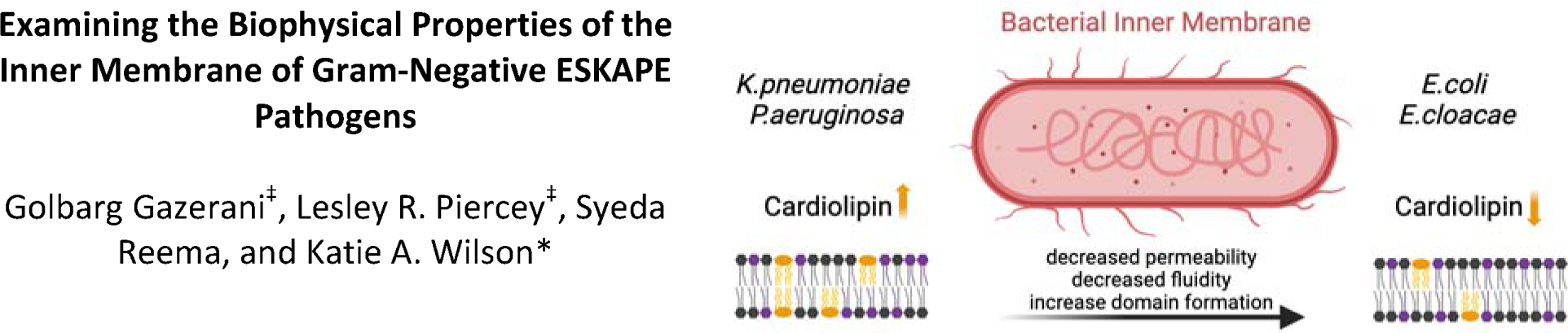
“For Table of Contents Use Only”.

