## Supporting Information for "Examining the Biophysical Properties of the Inner Membrane of Gram-Negative ESKAPE Pathogens"

[‡]These authors contributed equally

### Table of Contents

Table S1. Composition of the bacterial inner membranes ........................................................S2

Table S2. Headgroup composition of the bacterial inner membranes .......................................S2

Table S3. Average lipid unsaturation of the bacterial inner membranes.....................................S2

Figure S1. Membrane thickness and area per lipid over the triplicate 10 µs coarse grain molecular dynamics simulations ..................................................................................................S3

Figure S2. Average lipid order for each individual lipid species..................................................S4

Figure S3. Lipid order over the triplicate 10 µs coarse grain molecular dynamics simulations ....S3

Figure S4. Mean membrane curvature mapped to the surface of the upper and lower leaflets .S5

Figure S5. Depletion enrichment index in the *E.cloacae* inner membrane .................................S6

Figure S6. Depletion enrichment index in the *P.aeruginosa* inner membrane ............................S7

Figure S7. Depletion enrichment index in the *K.pneumonia* inner membrane ...........................S8

Figure S8. Depletion enrichment index in the *E.coli* inner membrane .......................................S9

**Table S1.** Composition of the bacterial inner membranes studied in the current work. The lipid nomenclature for PG and PE lipids follows standard Martini conventions, where the first two letters of a phospholipid name indicate the lipid tail groups and the last two letters indicate the lipid headgroup. Since cardiolipin contains 4 lipid tails, the first 4 letters of the lipid name indicate the cardiolipin tails. The lipid tails included in the present work include 16:0 (P), 16:1 (O), 18:1 (V), cyclic 17 (M) and cyclic 19 (N).

| | *E.coli* | *E.cloacae* | *K.pneumoniae* | *P.aeruginosa* |
|---|---|---|---|---|
| OPMN-CL | 1 | 1 | 1 | 2 |
| OPMV-CL | 1 | 1 | 1 | 3 |
| MPPV-CL | 2 | 1 | 2 | 5 |
| MPPM-CL | 1 | 0 | 3 | 2 |
| PMPG | 7 | 8 | 3 | 0 |
| PNPG | 2 | 0 | 1 | 5 |
| OVPG | 1 | 0 | 0 | 6 |
| PVPG | 3 | 0 | 0 | 12 |
| POPG | 2 | 3 | 1 | 0 |
| MOPG | 0 | 10 | 0 | 0 |
| PMPE | 17 | 20 | 32 | 3 |
| OVPE | 16 | 7 | 10 | 2 |
| PNPE | 6 | 10 | 18 | 8 |
| MVPE | 13 | 23 | 13 | 21 |
| PVPE | 28 | 6 | 0 | 17 |
| MOPE | 0 | 7 | 0 | 0 |
| NVPE | 0 | 0 | 0 | 5 |
| DVPE | 0 | 0 | 0 | 9 |
| DPPE | 0 | 3 | 15 | 0 |

**Table S2.** Headgroup composition of the bacterial inner membranes simulated in the current work.

| | *E.coli* | *E.cloacae* | *K.pneumoniae* | *P.aeruginosa* |
|---|---|---|---|---|
| CL | 5 | 3 | 7 | 12 |
| PG | 15 | 21 | 5 | 23 |
| PE | 80 | 76 | 88 | 65 |

**Table S3.** Average lipid unsaturation of the bacterial inner membranes simulated in the current work.

| | *E.coli* | *E. cloacae* | *K. pneumoniae* | *P. aeruginosa* |
|---|---|---|---|---|
| Unsaturations per lipid tail | 0.41 | 0.32 | 0.18 | 0.46 |
| Cyclic groups per lipid tail | 0.25 | 0.40 | 0.36 | 0.26 |

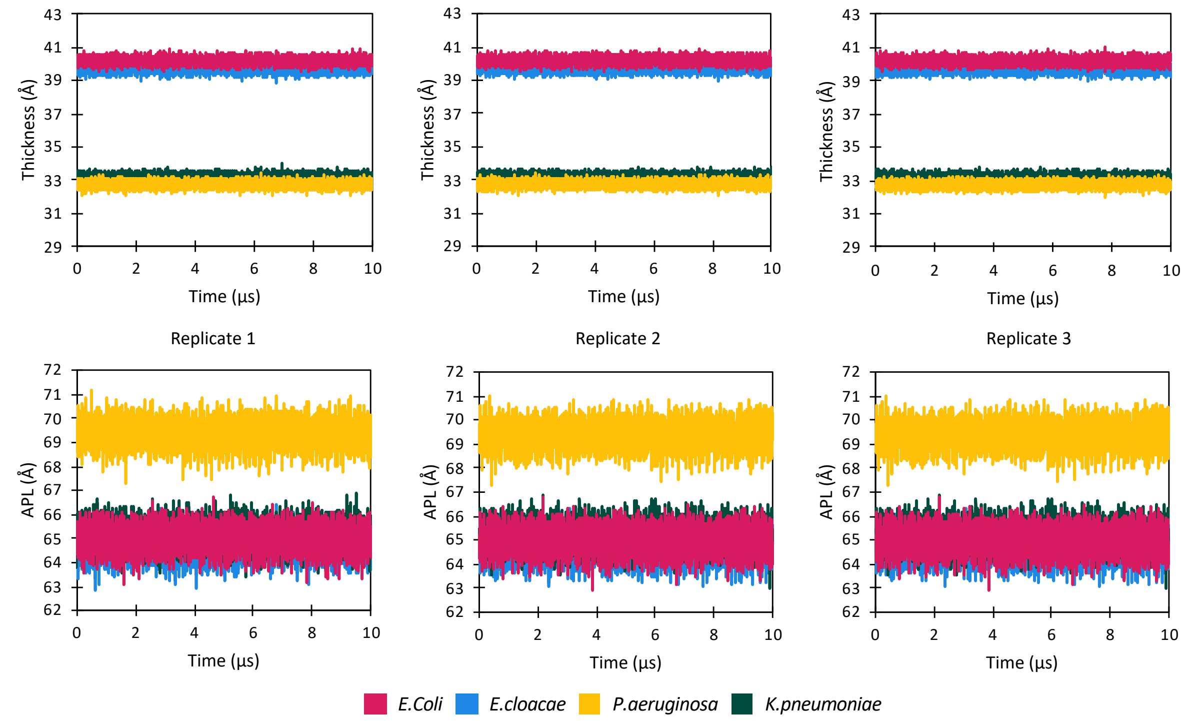


**Figure S1.** Membrane thickness (top) and area per lipid (bottom) over the triplicate 10 µs coarse grain molecular dynamics simulations for the *E.coli, E.cloacae*, *K.pneumoniae* and *P.aeruginosa* inner membrane.

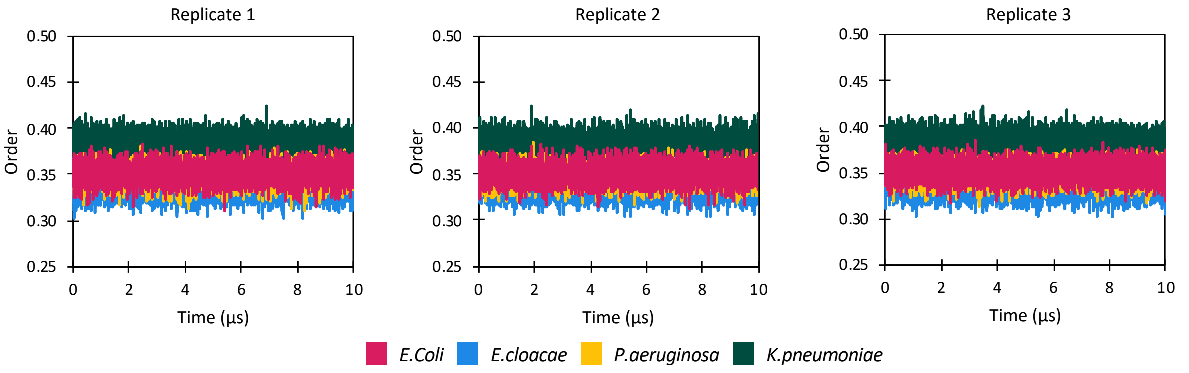


**Figure S2.** Lipid order over the triplicate 10 µs coarse grain molecular dynamics simulations for the *E.coli, E.cloacae*, *K.pneumoniae* and *P.aeruginosa* inner membrane.

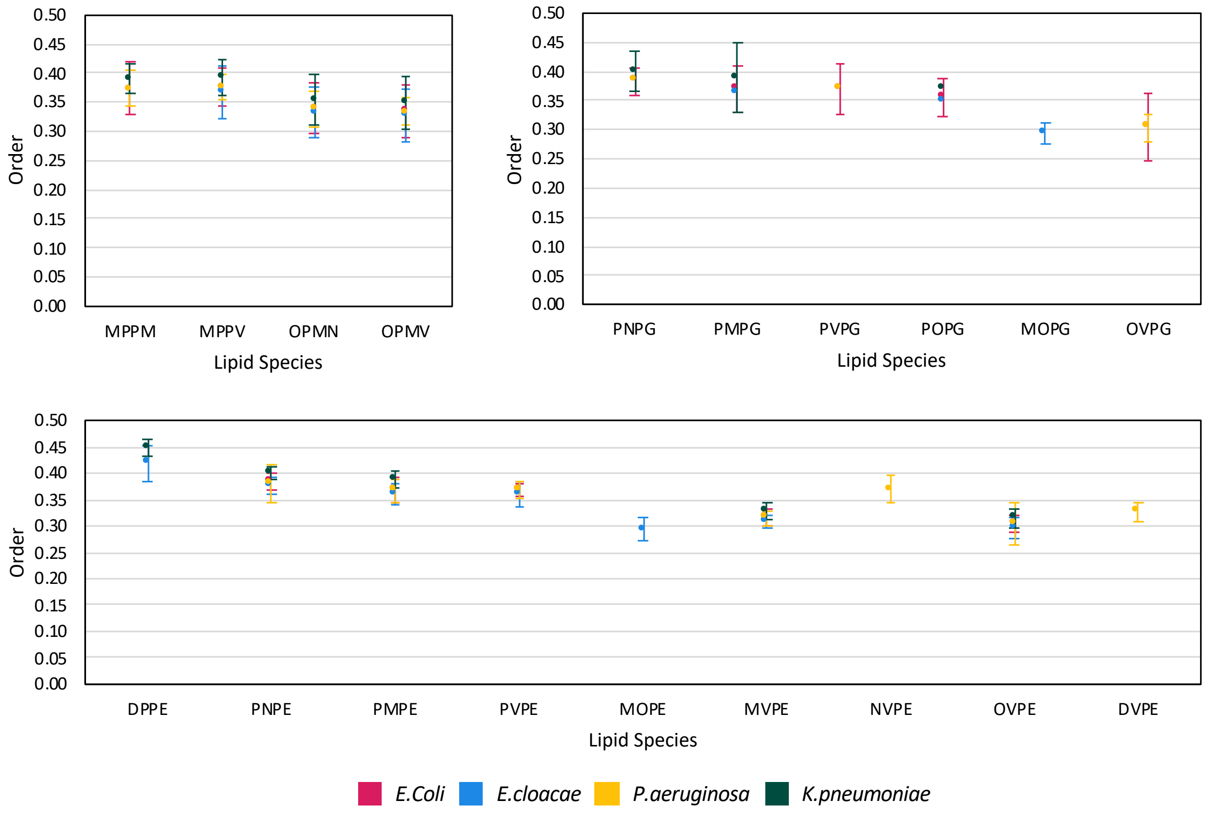


**Figure S3.** Average lipid order for each individual lipid species during triplicate 10 µs coarse grain molecular dynamics simulations of the *E.coli, E.cloacae*, *K.pneumoniae* and *P.aeruginosa* inner membrane. Reported errors are the standard deviation across the three replicates.

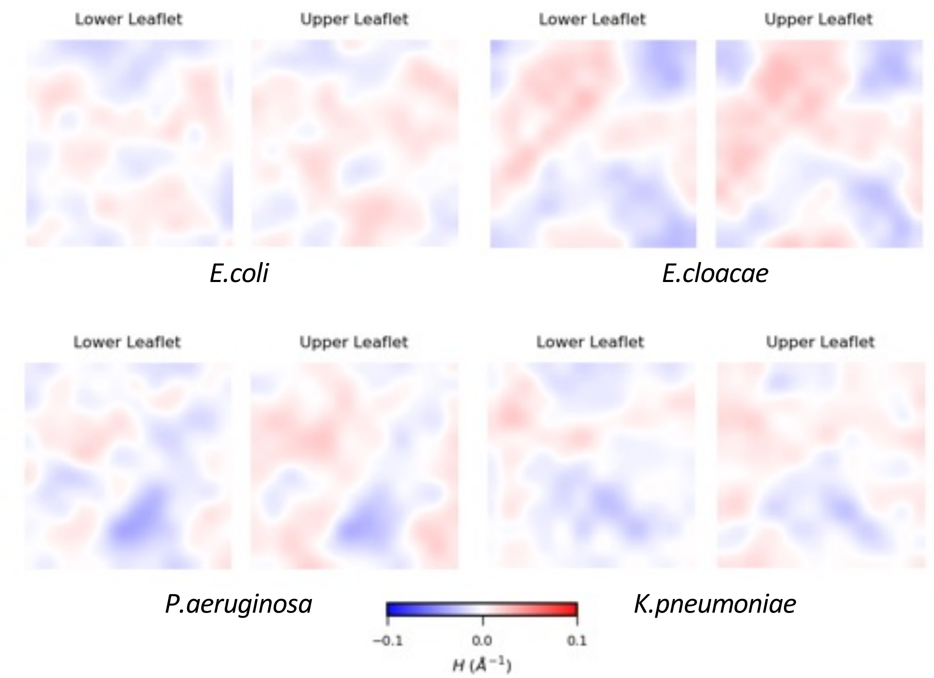


**Figure S4.** Mean membrane curvature (H) mapped to the surface of the upper and lower leaflets of the *E.coli, E.cloacae*, *K.pneumoniae* and *P.aeruginosa* inner membranes during triplicate 10 µs coarse grain molecular dynamics simulations.

a)

| | DPPE | MOPE | MVPE | OVPE | PMPE | PNPE | PVPE | MOPG | PMPG | POPG | MPPV | OPMN | OPMV |
|---|---|---|---|---|---|---|---|---|---|---|---|---|---|
| DPPE | 1.0 | 1.0 | 1.0 | 1.0 | 1.0 | 1.0 | 1.0 | 1.0 | 1.1 | 1.0 | 1.0 | 1.1 | 1.0 |
| MOPE | 1.0 | 1.0 | 1.0 | 1.0 | 1.0 | 1.0 | 1.0 | 1.0 | 1.0 | 1.0 | 1.0 | 1.0 | 1.0 |
| MVPE | 1.0 | 1.0 | 1.0 | 1.0 | 1.0 | 1.0 | 1.0 | 1.0 | 1.0 | 1.0 | 1.0 | 1.0 | 1.1 |
| OVPE | 0.9 | 1.0 | 1.0 | 1.0 | 0.9 | 1.0 | 1.0 | 1.0 | 1.0 | 1.0 | 1.0 | 1.1 | 1.0 |
| PMPE | 1.0 | 1.0 | 1.0 | 1.0 | 1.0 | 1.0 | 1.0 | 1.0 | 1.0 | 1.0 | 1.1 | 1.0 | 1.0 |
| PNPE | 1.0 | 1.0 | 1.0 | 1.0 | 1.0 | 1.0 | 1.0 | 1.0 | 1.1 | 1.0 | 1.1 | 1.0 | 1.0 |
| PVPE | 1.0 | 1.0 | 1.0 | 1.0 | 1.0 | 1.0 | 1.0 | 1.0 | 1.0 | 1.0 | 1.1 | 1.0 | 1.1 |
| MOPG | 1.0 | 1.0 | 1.0 | 1.0 | 1.0 | 1.0 | 1.0 | 0.9 | 0.9 | 0.9 | 0.8 | 0.8 | 0.8 |
| PMPG | 1.0 | 1.0 | 1.0 | 1.0 | 1.0 | 1.0 | 1.0 | 0.9 | 0.9 | 0.9 | 0.9 | 0.9 | 0.9 |
| POPG | 1.0 | 1.0 | 1.0 | 1.0 | 1.0 | 1.0 | 1.0 | 0.9 | 0.9 | 0.9 | 0.9 | 0.8 | 0.9 |
| MPPV | 1.3 | 1.3 | 1.4 | 1.4 | 1.4 | 1.4 | 1.4 | 1.1 | 1.1 | 1.2 | 0.9 | 1.0 | 1.0 |
| OPMN | 1.4 | 1.4 | 1.3 | 1.4 | 1.3 | 1.3 | 1.3 | 1.1 | 1.1 | 1.1 | 1.0 | 0.7 | 1.0 |
| OPMV | 1.3 | 1.3 | 1.4 | 1.3 | 1.3 | 1.3 | 1.4 | 1.1 | 1.1 | 1.2 | 1.0 | 1.0 | 0.8 |

b)

| | DPPE | MOPE | MVPE | OVPE | PMPE | PNPE | PVPE | MOPG | PMPG | POPG | MPPV | OPMN | OPMV |
|---|---|---|---|---|---|---|---|---|---|---|---|---|---|
| DPPE | 1.0 | 1.0 | 1.0 | 1.0 | 1.0 | 1.0 | 1.0 | 1.0 | 1.1 | 1.0 | 1.1 | 1.0 | 1.0 |
| MOPE | 1.0 | 1.0 | 1.0 | 1.0 | 1.0 | 0.9 | 1.0 | 1.0 | 1.0 | 1.0 | 1.0 | 1.0 | 1.0 |
| MVPE | 1.0 | 1.0 | 1.0 | 1.0 | 1.0 | 1.0 | 1.0 | 1.0 | 1.0 | 1.0 | 1.0 | 1.0 | 1.1 |
| OVPE | 0.9 | 1.0 | 1.0 | 1.0 | 1.0 | 0.9 | 1.0 | 1.0 | 1.0 | 1.0 | 1.0 | 1.0 | 1.0 |
| PMPE | 1.0 | 1.0 | 1.0 | 1.0 | 1.0 | 1.0 | 1.0 | 1.0 | 1.0 | 1.0 | 1.1 | 1.0 | 1.0 |
| PNPE | 1.0 | 1.0 | 1.0 | 1.0 | 1.0 | 1.0 | 1.0 | 1.0 | 1.1 | 1.0 | 1.1 | 1.1 | 1.0 |
| PVPE | 1.0 | 1.0 | 1.0 | 1.0 | 1.0 | 1.0 | 1.0 | 1.0 | 1.0 | 1.0 | 1.0 | 1.0 | 1.0 |
| MOPG | 1.0 | 1.0 | 1.0 | 1.0 | 1.0 | 1.0 | 1.0 | 0.9 | 0.9 | 0.9 | 0.9 | 0.8 | 0.8 |
| PMPG | 1.0 | 1.0 | 1.0 | 1.0 | 1.0 | 1.0 | 1.0 | 0.9 | 0.9 | 0.9 | 0.8 | 0.9 | 0.9 |
| POPG | 1.0 | 1.0 | 1.0 | 1.0 | 1.0 | 1.0 | 1.0 | 0.9 | 0.9 | 0.9 | 0.8 | 0.9 | 0.9 |
| MPPV | 1.4 | 1.3 | 1.4 | 1.4 | 1.4 | 1.4 | 1.3 | 1.1 | 1.1 | 1.1 | 0.9 | 1.1 | 0.8 |
| OPMN | 1.3 | 1.3 | 1.3 | 1.3 | 1.3 | 1.4 | 1.3 | 1.1 | 1.1 | 1.1 | 1.1 | 0.9 | 1.1 |
| OPMV | 1.3 | 1.4 | 1.4 | 1.3 | 1.3 | 1.3 | 1.3 | 1.1 | 1.1 | 1.1 | 0.8 | 1.1 | 0.8 |

c)

| | DPPE | MOPE | MVPE | OVPE | PMPE | PNPE | PVPE | MOPG | PMPG | POPG | MPPV | OPMN | OPMV |
|---|---|---|---|---|---|---|---|---|---|---|---|---|---|
| DPPE | 1.0 | 1.0 | 1.0 | 1.0 | 1.0 | 1.0 | 1.0 | 1.0 | 1.1 | 1.0 | 1.1 | 1.0 | 1.1 |
| MOPE | 1.0 | 1.0 | 1.0 | 1.0 | 1.0 | 1.0 | 1.0 | 1.0 | 1.0 | 1.0 | 1.0 | 1.0 | 1.0 |
| MVPE | 1.0 | 1.0 | 1.0 | 1.0 | 1.0 | 1.0 | 1.0 | 1.0 | 1.0 | 1.0 | 1.1 | 1.0 | 1.0 |
| OVPE | 0.9 | 1.0 | 1.0 | 1.0 | 1.0 | 1.0 | 1.0 | 1.0 | 1.0 | 1.0 | 1.0 | 1.0 | 1.1 |
| PMPE | 1.0 | 1.0 | 1.0 | 1.0 | 1.0 | 1.0 | 1.0 | 1.0 | 1.0 | 1.0 | 1.1 | 1.0 | 1.0 |
| PNPE | 1.0 | 1.0 | 1.0 | 1.0 | 1.0 | 1.0 | 1.0 | 1.0 | 1.0 | 1.0 | 1.1 | 1.1 | 1.0 |
| PVPE | 1.0 | 1.0 | 1.0 | 1.0 | 1.0 | 1.0 | 1.0 | 1.0 | 1.0 | 1.0 | 1.0 | 1.1 | 1.0 |
| MOPG | 1.0 | 1.0 | 1.0 | 1.0 | 1.0 | 1.0 | 1.0 | 0.9 | 0.9 | 0.9 | 0.8 | 0.8 | 0.8 |
| PMPG | 1.0 | 1.0 | 1.0 | 1.0 | 1.0 | 1.0 | 1.0 | 0.9 | 0.9 | 0.9 | 0.9 | 0.9 | 0.9 |
| POPG | 1.0 | 1.0 | 1.0 | 1.0 | 1.0 | 1.0 | 1.0 | 0.9 | 0.9 | 0.9 | 0.9 | 0.9 | 0.8 |
| MPPV | 1.4 | 1.3 | 1.4 | 1.3 | 1.4 | 1.4 | 1.3 | 1.1 | 1.1 | 1.1 | 0.8 | 1.1 | 1.1 |
| OPMN | 1.3 | 1.3 | 1.3 | 1.3 | 1.3 | 1.4 | 1.4 | 1.1 | 1.1 | 1.2 | 1.1 | 0.7 | 0.9 |
| OPMV | 1.3 | 1.4 | 1.4 | 1.4 | 1.3 | 1.3 | 1.3 | 1.1 | 1.1 | 1.0 | 1.0 | 0.9 | 0.8 |

0.6 1 1.4

**Figure S5.** Depletion enrichment index (DEI) reporting lipid clustering in the *E.cloacae* inner membrane the three replicates (a, b and c) 10 µs of coarse grain molecular dynamics simulation.

a)

| | DVPE | MVPE | NVPE | OVPE | PMPE | PNPE | PVPE | OVPG | PNPG | PVPG | MPPM | MPPV | OPMN | OPMV |
|---|---|---|---|---|---|---|---|---|---|---|---|---|---|---|
| DVPE | 1.0 | 1.0 | 0.9 | 1.0 | 0.9 | 0.9 | 1.0 | 1.1 | 1.0 | 1.0 | 1.0 | 1.0 | 1.0 | 1.0 |
| MVPE | 0.9 | 0.9 | 0.9 | 1.0 | 0.9 | 0.9 | 0.9 | 1.0 | 1.0 | 1.0 | 1.0 | 1.0 | 1.0 | 1.0 |
| NVPE | 0.9 | 0.9 | 1.0 | 0.9 | 1.0 | 1.0 | 1.0 | 1.0 | 1.0 | 1.0 | 1.0 | 1.0 | 1.0 | 1.1 |
| OVPE | 1.0 | 0.9 | 0.9 | 0.9 | 1.0 | 0.9 | 1.0 | 1.0 | 1.0 | 1.0 | 1.0 | 1.0 | 1.0 | 1.0 |
| PMPE | 0.9 | 0.9 | 1.0 | 1.0 | 0.9 | 1.0 | 0.9 | 1.0 | 1.0 | 1.0 | 1.0 | 1.0 | 1.1 | 1.0 |
| PNPE | 0.9 | 1.0 | 1.0 | 0.9 | 1.0 | 1.0 | 1.0 | 1.0 | 1.0 | 1.0 | 1.1 | 1.1 | 1.0 | 1.0 |
| PVPE | 1.0 | 0.9 | 1.0 | 1.0 | 0.9 | 1.0 | 0.9 | 1.0 | 1.0 | 1.0 | 1.0 | 1.0 | 1.0 | 1.0 |
| OVPG | 1.0 | 1.0 | 1.0 | 1.0 | 1.0 | 0.9 | 1.0 | 0.9 | 0.9 | 0.9 | 0.9 | 0.9 | 0.9 | 0.9 |
| PNPG | 1.0 | 1.0 | 1.0 | 1.0 | 1.0 | 1.0 | 1.0 | 0.9 | 0.9 | 0.9 | 0.9 | 0.9 | 0.9 | 0.9 |
| PVPG | 1.0 | 1.0 | 1.0 | 1.0 | 1.0 | 1.0 | 1.0 | 0.9 | 0.9 | 0.9 | 0.9 | 0.9 | 0.9 | 0.9 |
| MPPM | 1.2 | 1.3 | 1.3 | 1.2 | 1.3 | 1.4 | 1.3 | 1.1 | 1.2 | 1.2 | 1.1 | 1.2 | 1.1 | 1.1 |
| MPPV | 1.3 | 1.3 | 1.3 | 1.3 | 1.3 | 1.4 | 1.3 | 1.1 | 1.2 | 1.2 | 1.2 | 1.1 | 1.1 | 1.1 |
| OPMN | 1.3 | 1.3 | 1.3 | 1.3 | 1.4 | 1.3 | 1.3 | 1.2 | 1.2 | 1.2 | 1.1 | 1.1 | 1.0 | 1.1 |
| OPMV | 1.3 | 1.3 | 1.3 | 1.3 | 1.3 | 1.3 | 1.3 | 1.2 | 1.2 | 1.2 | 1.1 | 1.1 | 1.1 | 1.1 |

b)

| | DVPE | MVPE | NVPE | OVPE | PMPE | PNPE | PVPE | OVPG | PNPG | PVPG | MPPM | MPPV | OPMN | OPMV |
|---|---|---|---|---|---|---|---|---|---|---|---|---|---|---|
| DVPE | 1.0 | 1.0 | 0.9 | 1.0 | 0.9 | 0.9 | 1.0 | 1.0 | 1.0 | 1.0 | 1.0 | 1.0 | 1.0 | 1.0 |
| MVPE | 1.0 | 0.9 | 0.9 | 0.9 | 0.9 | 0.9 | 0.9 | 1.0 | 1.0 | 1.0 | 1.0 | 1.0 | 1.0 | 1.0 |
| NVPE | 0.9 | 0.9 | 0.9 | 1.0 | 1.0 | 1.0 | 1.0 | 1.0 | 1.0 | 1.0 | 1.1 | 1.0 | 1.0 | 1.0 |
| OVPE | 1.0 | 0.9 | 0.9 | 0.9 | 0.9 | 0.9 | 1.0 | 1.0 | 0.9 | 1.0 | 0.9 | 1.0 | 1.0 | 1.0 |
| PMPE | 0.9 | 0.9 | 1.0 | 0.9 | 0.9 | 1.0 | 0.9 | 1.0 | 1.0 | 1.0 | 1.1 | 1.0 | 1.1 | 1.0 |
| PNPE | 0.9 | 0.9 | 1.0 | 0.9 | 1.0 | 1.0 | 1.0 | 1.0 | 1.0 | 1.0 | 1.1 | 1.0 | 1.0 | 1.0 |
| PVPE | 1.0 | 1.0 | 1.0 | 1.0 | 0.9 | 1.0 | 0.9 | 1.0 | 1.0 | 1.0 | 1.0 | 1.0 | 1.0 | 1.0 |
| OVPG | 1.0 | 1.0 | 0.9 | 1.0 | 0.9 | 0.9 | 1.0 | 0.9 | 0.9 | 0.9 | 0.8 | 0.9 | 0.9 | 0.9 |
| PNPG | 1.0 | 1.0 | 1.0 | 0.9 | 1.0 | 1.0 | 1.0 | 0.9 | 0.9 | 0.9 | 0.9 | 0.9 | 0.9 | 0.9 |
| PVPG | 1.0 | 1.0 | 1.0 | 1.0 | 1.0 | 1.0 | 1.0 | 0.9 | 0.9 | 0.9 | 0.9 | 0.9 | 0.9 | 0.9 |
| MPPM | 1.3 | 1.3 | 1.3 | 1.2 | 1.4 | 1.4 | 1.3 | 1.1 | 1.2 | 1.2 | 1.1 | 1.2 | 1.1 | 1.0 |
| MPPV | 1.3 | 1.3 | 1.3 | 1.2 | 1.3 | 1.3 | 1.3 | 1.1 | 1.2 | 1.2 | 1.2 | 1.1 | 1.2 | 1.1 |
| OPMN | 1.3 | 1.3 | 1.3 | 1.3 | 1.4 | 1.3 | 1.3 | 1.2 | 1.2 | 1.1 | 1.1 | 1.1 | 1.0 | 1.1 |
| OPMV | 1.3 | 1.3 | 1.3 | 1.3 | 1.3 | 1.3 | 1.3 | 1.2 | 1.1 | 1.2 | 1.0 | 1.1 | 1.1 | 1.0 |

c)

| | DVPE | MVPE | NVPE | OVPE | PMPE | PNPE | PVPE | OVPG | PNPG | PVPG | MPPM | MPPV | OPMN | OPMV |
|---|---|---|---|---|---|---|---|---|---|---|---|---|---|---|
| DVPE | 1.0 | 1.0 | 0.9 | 1.0 | 0.9 | 0.9 | 1.0 | 1.0 | 1.0 | 1.0 | 1.0 | 1.0 | 1.0 | 1.0 |
| MVPE | 1.0 | 0.9 | 0.9 | 1.0 | 1.0 | 0.9 | 0.9 | 1.0 | 1.0 | 1.0 | 1.0 | 1.0 | 1.0 | 1.0 |
| NVPE | 0.9 | 1.0 | 0.9 | 0.9 | 1.0 | 1.0 | 0.9 | 1.0 | 1.0 | 1.0 | 1.1 | 1.1 | 1.0 | 1.0 |
| OVPE | 0.9 | 1.0 | 0.9 | 0.8 | 0.9 | 0.9 | 0.9 | 1.0 | 1.0 | 1.0 | 0.9 | 1.0 | 1.0 | 1.0 |
| PMPE | 0.9 | 1.0 | 1.0 | 0.9 | 1.0 | 1.0 | 0.9 | 1.0 | 1.0 | 1.0 | 1.1 | 1.0 | 1.0 | 1.0 |
| PNPE | 0.9 | 0.9 | 1.0 | 0.9 | 1.0 | 0.9 | 1.0 | 1.0 | 1.0 | 1.0 | 1.1 | 1.1 | 1.1 | 1.0 |
| PVPE | 1.0 | 1.0 | 0.9 | 1.0 | 0.9 | 1.0 | 1.0 | 1.0 | 1.0 | 1.0 | 1.0 | 1.0 | 1.0 | 1.0 |
| OVPG | 1.0 | 1.0 | 1.0 | 1.0 | 1.0 | 0.9 | 1.0 | 1.0 | 0.9 | 0.9 | 0.8 | 0.9 | 0.9 | 0.9 |
| PNPG | 1.0 | 1.0 | 1.0 | 1.0 | 1.0 | 1.0 | 1.0 | 0.9 | 0.9 | 0.9 | 0.9 | 0.9 | 1.0 | 0.9 |
| PVPG | 1.0 | 1.0 | 1.0 | 1.0 | 1.0 | 1.0 | 1.0 | 0.9 | 0.9 | 0.9 | 0.9 | 0.9 | 0.9 | 0.9 |
| MPPM | 1.3 | 1.3 | 1.4 | 1.2 | 1.4 | 1.4 | 1.3 | 1.1 | 1.2 | 1.2 | 1.0 | 1.1 | 1.2 | 1.1 |
| MPPV | 1.3 | 1.3 | 1.4 | 1.3 | 1.3 | 1.4 | 1.3 | 1.2 | 1.2 | 1.2 | 1.1 | 1.1 | 1.1 | 1.1 |
| OPMN | 1.3 | 1.3 | 1.3 | 1.3 | 1.3 | 1.3 | 1.3 | 1.1 | 1.2 | 1.2 | 1.2 | 1.1 | 0.9 | 1.0 |
| OPMV | 1.3 | 1.3 | 1.3 | 1.3 | 1.3 | 1.3 | 1.3 | 1.2 | 1.1 | 1.2 | 1.1 | 1.1 | 1.0 | 1.1 |

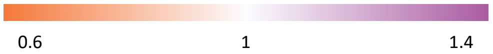


**Figure S6.** Depletion enrichment index (DEI) reporting lipid clustering in the *P.aeruginosa* inner membrane the three replicates (a, b and c) 10 µs of coarse grain molecular dynamics simulation.

a)

| | DPPE | MVPE | OVPE | PMPE | PNPE | PMPG | PNPG | POPG | MPPM | MPPV | OPMN | OPMV |
|---|---|---|---|---|---|---|---|---|---|---|---|---|
| DPPE | 1.0 | 1.0 | 1.0 | 1.0 | 1.0 | 1.0 | 1.0 | 1.0 | 1.0 | 1.0 | 1.0 | 1.0 |
| MVPE | 1.0 | 1.0 | 1.0 | 1.0 | 1.0 | 1.0 | 1.0 | 1.0 | 1.0 | 1.0 | 1.0 | 1.0 |
| OVPE | 0.9 | 1.0 | 1.0 | 0.9 | 0.9 | 1.0 | 1.0 | 1.0 | 1.0 | 1.0 | 1.0 | 1.1 |
| PMPE | 1.0 | 1.0 | 1.0 | 1.0 | 1.0 | 1.0 | 1.0 | 1.0 | 1.0 | 1.0 | 1.0 | 1.0 |
| PNPE | 1.0 | 1.0 | 1.0 | 1.0 | 1.0 | 1.0 | 1.0 | 1.0 | 1.0 | 1.0 | 1.0 | 1.0 |
| PMPG | 1.0 | 1.0 | 1.0 | 1.0 | 1.0 | 0.8 | 0.8 | 0.8 | 0.8 | 0.8 | 0.8 | 0.8 |
| PNPG | 1.0 | 1.0 | 1.0 | 1.0 | 1.0 | 0.8 | 0.8 | 0.8 | 0.8 | 0.8 | 0.8 | 0.8 |
| POPG | 1.0 | 1.0 | 1.0 | 1.0 | 1.0 | 0.8 | 0.8 | 0.7 | 0.8 | 0.8 | 0.8 | 0.7 |
| MPPM | 1.4 | 1.3 | 1.3 | 1.3 | 1.3 | 1.0 | 1.0 | 1.1 | 0.9 | 0.9 | 1.0 | 0.9 |
| MPPV | 1.3 | 1.3 | 1.4 | 1.3 | 1.3 | 1.0 | 1.0 | 1.1 | 0.9 | 0.8 | 0.9 | 0.9 |
| OPMN | 1.3 | 1.3 | 1.4 | 1.3 | 1.3 | 1.0 | 1.1 | 1.0 | 0.9 | 0.9 | 0.7 | 0.9 |
| OPMV | 1.3 | 1.3 | 1.4 | 1.3 | 1.3 | 1.0 | 1.0 | 1.0 | 0.9 | 0.9 | 0.9 | 0.8 |

b)

| | DPPE | MVPE | OVPE | PMPE | PNPE | PMPG | PNPG | POPG | MPPM | MPPV | OPMN | OPMV |
|---|---|---|---|---|---|---|---|---|---|---|---|---|
| DPPE | 1.0 | 1.0 | 1.0 | 1.0 | 1.0 | 1.0 | 1.0 | 1.0 | 1.1 | 1.0 | 1.0 | 1.0 |
| MVPE | 1.0 | 1.0 | 1.0 | 1.0 | 1.0 | 1.0 | 1.0 | 1.0 | 1.0 | 1.0 | 1.0 | 1.1 |
| OVPE | 0.9 | 1.0 | 1.0 | 0.9 | 0.9 | 1.0 | 1.0 | 1.0 | 1.0 | 1.0 | 1.0 | 1.1 |
| PMPE | 1.0 | 1.0 | 1.0 | 1.0 | 1.0 | 1.0 | 1.0 | 1.0 | 1.0 | 1.0 | 1.0 | 1.0 |
| PNPE | 1.0 | 1.0 | 1.0 | 1.0 | 1.0 | 1.0 | 1.0 | 1.0 | 1.0 | 1.0 | 1.0 | 1.0 |
| PMPG | 1.0 | 1.0 | 1.0 | 1.0 | 1.0 | 0.8 | 0.9 | 0.9 | 0.8 | 0.8 | 0.8 | 0.8 |
| PNPG | 1.0 | 1.0 | 1.0 | 1.0 | 1.0 | 0.9 | 0.7 | 0.8 | 0.8 | 0.8 | 0.8 | 0.9 |
| POPG | 1.0 | 1.0 | 1.0 | 1.0 | 1.0 | 0.9 | 0.8 | 0.7 | 0.8 | 0.8 | 0.7 | 0.9 |
| MPPM | 1.4 | 1.3 | 1.3 | 1.3 | 1.4 | 1.0 | 1.0 | 1.1 | 0.9 | 0.9 | 0.9 | 0.9 |
| MPPV | 1.3 | 1.3 | 1.3 | 1.3 | 1.4 | 1.0 | 1.0 | 1.0 | 0.9 | 0.9 | 0.9 | 0.9 |
| OPMN | 1.3 | 1.3 | 1.3 | 1.3 | 1.3 | 1.0 | 1.0 | 0.9 | 0.8 | 0.9 | 0.8 | 0.9 |
| OPMV | 1.3 | 1.4 | 1.4 | 1.3 | 1.3 | 1.1 | 1.1 | 1.2 | 0.8 | 0.9 | 0.9 | 0.7 |

c)

| | DPPE | MVPE | OVPE | PMPE | PNPE | PMPG | PNPG | POPG | MPPM | MPPV | OPMN | OPMV |
|---|---|---|---|---|---|---|---|---|---|---|---|---|
| DPPE | 1.0 | 1.0 | 1.0 | 1.0 | 1.0 | 1.0 | 1.0 | 1.0 | 1.0 | 1.1 | 1.0 | 1.0 |
| MVPE | 1.0 | 1.0 | 1.0 | 1.0 | 1.0 | 1.0 | 1.0 | 1.0 | 1.0 | 1.0 | 1.0 | 1.0 |
| OVPE | 0.9 | 1.0 | 1.0 | 1.0 | 0.9 | 1.0 | 1.0 | 1.0 | 1.0 | 1.0 | 1.0 | 1.1 |
| PMPE | 1.0 | 1.0 | 1.0 | 1.0 | 1.0 | 1.0 | 1.0 | 1.0 | 1.0 | 1.0 | 1.0 | 1.0 |
| PNPE | 1.0 | 1.0 | 1.0 | 1.0 | 1.0 | 1.0 | 1.0 | 1.0 | 1.0 | 1.0 | 1.0 | 1.0 |
| PMPG | 1.0 | 1.0 | 1.0 | 1.0 | 1.0 | 0.8 | 0.9 | 0.9 | 0.8 | 0.8 | 0.8 | 0.8 |
| PNPG | 1.0 | 1.0 | 1.0 | 1.0 | 1.0 | 0.9 | 0.7 | 0.8 | 0.8 | 0.8 | 0.8 | 0.7 |
| POPG | 1.0 | 1.0 | 1.0 | 1.0 | 1.0 | 0.9 | 0.8 | 0.7 | 0.7 | 0.8 | 0.8 | 0.8 |
| MPPM | 1.3 | 1.3 | 1.3 | 1.3 | 1.3 | 1.0 | 1.0 | 1.0 | 0.9 | 0.9 | 1.0 | 0.9 |
| MPPV | 1.4 | 1.3 | 1.3 | 1.3 | 1.3 | 1.0 | 1.1 | 1.0 | 0.9 | 0.9 | 0.9 | 0.9 |
| OPMN | 1.3 | 1.4 | 1.3 | 1.3 | 1.3 | 1.0 | 1.0 | 1.0 | 0.9 | 0.9 | 0.7 | 0.8 |
| OPMV | 1.3 | 1.3 | 1.4 | 1.3 | 1.3 | 1.0 | 1.0 | 1.1 | 0.9 | 0.8 | 0.9 | 0.8 |

0.6 1 1.4

**Figure S7.** Depletion enrichment index (DEI) reporting lipid clustering in the K.pneumonia inner membrane the three replicates (a, b and c) 10 µs of coarse grain molecular dynamics simulation.

a)

| | MVPE | OVPE | PMPE | PNPE | PVPE | OVPG | PMPG | PNPG | POPG | PVPG | MPPM | MPPV | OPMN | OPMV |
|---|---|---|---|---|---|---|---|---|---|---|---|---|---|---|
| MVPE | 1.0 | 1.0 | 1.0 | 1.0 | 1.0 | 1.0 | 1.0 | 1.0 | 1.0 | 1.0 | 1.0 | 1.0 | 1.0 | 1.1 |
| OVPE | 1.0 | 1.0 | 0.9 | 0.9 | 1.0 | 1.1 | 1.0 | 1.0 | 1.0 | 1.0 | 1.0 | 1.0 | 1.0 | 1.0 |
| PMPE | 1.0 | 1.0 | 1.0 | 1.0 | 1.0 | 1.0 | 1.0 | 1.0 | 1.0 | 1.0 | 1.1 | 1.0 | 1.0 | 1.0 |
| PNPE | 1.0 | 1.0 | 1.0 | 1.0 | 1.0 | 1.0 | 1.0 | 1.1 | 1.0 | 1.0 | 1.1 | 1.0 | 1.1 | 1.0 |
| PVPE | 1.0 | 1.0 | 1.0 | 1.0 | 1.0 | 1.0 | 1.0 | 1.0 | 1.0 | 1.0 | 1.0 | 1.0 | 1.0 | 1.0 |
| OVPG | 1.0 | 1.1 | 1.0 | 0.9 | 1.0 | 0.7 | 0.8 | 0.9 | 0.9 | 0.8 | 0.8 | 0.8 | 0.9 | 0.9 |
| PMPG | 1.0 | 1.0 | 1.0 | 1.0 | 1.0 | 0.9 | 0.9 | 0.9 | 0.9 | 0.9 | 0.8 | 0.8 | 0.8 | 0.8 |
| PNPG | 1.0 | 1.0 | 1.0 | 1.1 | 1.0 | 0.9 | 0.9 | 0.9 | 1.0 | 0.9 | 0.9 | 0.9 | 0.8 | 0.8 |
| POPG | 1.0 | 1.0 | 1.0 | 1.0 | 1.0 | 1.0 | 0.9 | 0.9 | 0.9 | 0.9 | 0.8 | 0.8 | 0.8 | 0.9 |
| PVPG | 1.0 | 1.0 | 1.0 | 1.0 | 1.0 | 0.9 | 0.9 | 0.9 | 0.9 | 0.8 | 0.8 | 0.8 | 0.8 | 0.8 |
| MPPM | 1.3 | 1.3 | 1.4 | 1.4 | 1.3 | 1.0 | 1.1 | 1.1 | 1.0 | 1.1 | 1.0 | 1.0 | 0.9 | 1.0 |
| MPPV | 1.3 | 1.3 | 1.4 | 1.4 | 1.4 | 1.1 | 1.1 | 1.1 | 1.1 | 1.1 | 1.0 | 0.9 | 1.1 | 1.0 |
| OPMN | 1.3 | 1.3 | 1.4 | 1.4 | 1.3 | 1.1 | 1.1 | 1.1 | 1.1 | 1.1 | 0.9 | 1.1 | 0.9 | 0.9 |
| OPMV | 1.4 | 1.3 | 1.3 | 1.3 | 1.3 | 1.2 | 1.0 | 1.1 | 1.1 | 1.1 | 1.0 | 1.0 | 0.9 | 0.8 |

b)

| | MVPE | OVPE | PMPE | PNPE | PVPE | OVPG | PMPG | PNPG | POPG | PVPG | MPPM | MPPV | OPMN | OPMV |
|---|---|---|---|---|---|---|---|---|---|---|---|---|---|---|
| MVPE | 1.0 | 1.0 | 1.0 | 1.0 | 1.0 | 1.0 | 1.0 | 1.0 | 1.0 | 1.0 | 1.0 | 1.0 | 1.0 | 1.0 |
| OVPE | 1.0 | 1.0 | 0.9 | 0.9 | 1.0 | 1.1 | 1.0 | 1.0 | 1.0 | 1.0 | 1.0 | 1.0 | 1.0 | 1.0 |
| PMPE | 1.0 | 0.9 | 1.0 | 1.0 | 1.0 | 1.0 | 1.0 | 1.0 | 1.0 | 1.0 | 1.1 | 1.0 | 1.1 | 1.0 |
| PNPE | 1.0 | 1.0 | 1.0 | 1.0 | 1.0 | 1.0 | 1.0 | 1.0 | 1.0 | 1.0 | 1.1 | 1.0 | 1.1 | 1.0 |
| PVPE | 1.0 | 1.0 | 1.0 | 1.0 | 1.0 | 1.0 | 1.0 | 1.0 | 1.0 | 1.0 | 1.0 | 1.0 | 1.0 | 1.0 |
| OVPG | 1.0 | 1.1 | 1.0 | 1.0 | 1.0 | 0.6 | 0.9 | 0.8 | 0.9 | 0.8 | 0.8 | 0.8 | 0.8 | 0.9 |
| PMPG | 1.0 | 1.0 | 1.0 | 1.0 | 1.0 | 0.9 | 0.9 | 0.9 | 0.9 | 0.9 | 0.8 | 0.8 | 0.8 | 0.8 |
| PNPG | 1.0 | 1.0 | 1.0 | 1.0 | 1.0 | 0.8 | 0.9 | 0.8 | 0.9 | 0.9 | 0.9 | 0.8 | 0.9 | 0.9 |
| POPG | 1.0 | 1.0 | 1.0 | 1.0 | 1.0 | 0.9 | 0.9 | 0.9 | 0.8 | 0.9 | 0.8 | 0.9 | 0.8 | 0.8 |
| PVPG | 1.0 | 1.0 | 1.0 | 1.0 | 1.0 | 0.9 | 0.9 | 0.9 | 0.9 | 0.8 | 0.8 | 0.8 | 0.8 | 0.9 |
| MPPM | 1.3 | 1.3 | 1.4 | 1.4 | 1.3 | 1.0 | 1.1 | 1.1 | 1.0 | 1.1 | 0.9 | 1.0 | 1.1 | 0.9 |
| MPPV | 1.4 | 1.3 | 1.4 | 1.4 | 1.4 | 1.1 | 1.1 | 1.1 | 1.1 | 1.1 | 1.0 | 0.9 | 1.0 | 1.0 |
| OPMN | 1.3 | 1.3 | 1.4 | 1.4 | 1.3 | 1.1 | 1.1 | 1.1 | 1.0 | 1.1 | 1.1 | 1.0 | 0.8 | 1.0 |
| OPMV | 1.3 | 1.4 | 1.3 | 1.3 | 1.3 | 1.2 | 1.1 | 1.2 | 1.1 | 1.1 | 0.9 | 1.0 | 1.0 | 0.7 |

c)

| | MVPE | OVPE | PMPE | PNPE | PVPE | OVPG | PMPG | PNPG | POPG | PVPG | MPPM | MPPV | OPMN | OPMV |
|---|---|---|---|---|---|---|---|---|---|---|---|---|---|---|
| MVPE | 1.0 | 1.0 | 1.0 | 1.0 | 1.0 | 1.0 | 1.0 | 1.0 | 1.0 | 1.0 | 1.0 | 1.0 | 1.0 | 1.0 |
| OVPE | 1.0 | 1.0 | 0.9 | 0.9 | 1.0 | 1.1 | 1.0 | 1.0 | 1.0 | 1.0 | 1.0 | 1.0 | 1.0 | 1.0 |
| PMPE | 1.0 | 0.9 | 1.0 | 1.0 | 1.0 | 1.0 | 1.0 | 1.0 | 1.0 | 1.0 | 1.1 | 1.1 | 1.0 | 1.0 |
| PNPE | 1.0 | 1.0 | 1.0 | 1.0 | 1.0 | 1.0 | 1.0 | 1.0 | 1.0 | 1.0 | 1.1 | 1.0 | 1.1 | 1.0 |
| PVPE | 1.0 | 1.0 | 1.0 | 1.0 | 1.0 | 1.0 | 1.0 | 1.0 | 1.0 | 1.0 | 1.0 | 1.0 | 1.0 | 1.0 |
| OVPG | 1.0 | 1.1 | 1.0 | 1.0 | 1.0 | 0.7 | 0.8 | 0.8 | 0.9 | 0.9 | 0.8 | 0.8 | 0.7 | 0.8 |
| PMPG | 1.0 | 1.0 | 1.0 | 1.0 | 1.0 | 0.8 | 0.9 | 0.9 | 0.9 | 0.9 | 0.8 | 0.8 | 0.9 | 0.8 |
| PNPG | 1.0 | 1.0 | 1.0 | 1.0 | 1.0 | 0.9 | 0.9 | 0.8 | 0.9 | 0.9 | 0.9 | 0.9 | 0.9 | 0.8 |
| POPG | 1.0 | 1.0 | 1.0 | 1.0 | 1.0 | 0.9 | 0.9 | 0.9 | 0.9 | 0.9 | 0.9 | 0.8 | 0.8 | 0.9 |
| PVPG | 1.0 | 1.0 | 1.0 | 1.0 | 1.0 | 0.9 | 0.9 | 0.9 | 0.9 | 0.8 | 0.8 | 0.8 | 0.8 | 0.8 |
| MPPM | 1.4 | 1.3 | 1.4 | 1.4 | 1.3 | 1.1 | 1.1 | 1.2 | 1.1 | 1.1 | 0.7 | 0.9 | 1.0 | 1.0 |
| MPPV | 1.3 | 1.3 | 1.4 | 1.3 | 1.4 | 1.1 | 1.1 | 1.1 | 1.1 | 1.1 | 0.9 | 0.9 | 1.0 | 1.0 |
| OPMN | 1.3 | 1.3 | 1.3 | 1.4 | 1.3 | 0.9 | 1.1 | 1.1 | 1.1 | 1.0 | 1.0 | 1.0 | 0.8 | 1.0 |
| OPMV | 1.3 | 1.4 | 1.3 | 1.3 | 1.3 | 1.1 | 1.0 | 1.0 | 1.1 | 1.1 | 1.0 | 1.0 | 1.0 | 1.0 |

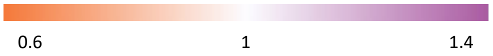


**Figure S8.** Depletion enrichment index (DEI) reporting lipid clustering in the E.coli inner membrane the three replicates (a, b and c) 10 μs of coarse grain molecular dynamics simulation.
